# Cytoophidium complexes resonate with cell fates

**DOI:** 10.1101/2024.09.20.614056

**Authors:** Yi-Lan Li, Ji-Long Liu

## Abstract

Metabolism is a fundamental characteristic of life. In 2010, we discovered that the metabolic enzyme CTP synthase (CTPS) can assemble a snake like structure inside cells, which we call the cytoophidium. Including CTPS, an increasing number of metabolic enzymes have been found to form cytoophidia in cells. However, the distribution and relationship among cytoophidia formed by different metabolic enzymes remain elusive. Here we investigate five metabolic enzymes that can form cytoophidia, namely Asn1, Bna5, CTPS (ie. Ura7), Glt1, and Prs5 in *Saccharomyces cerevisiae*. We find that multiple cytoophidia can be assembled into cytoophidium complexes by docking one after another. Glt1 cytoophidia tend to assemble in non-quiescent cells, while CTPS cytoophidia are more abundant in quiescent cells and form complexes with Prs5 and Asn1 cytoophidia. Blocking CTPS cytoophidium assembly can lead to a non-quiescent phenotype and increase the assembly of Glt1 cytoophidia, Bna5 cytoophidia, and cytoophidium complexes. Blocking CTPS cytoophidium assembly also inhibits the NAD biosynthesis pathway, which includes Bna5 and Sir2. Consistent with this result, the non-quiescent phenotype caused by blocking CTPS cytoophidium assembly can be rescued by blocking Glt1 cytoophidium assembly, supplementing nicotinic acid, or overexpressing Sir2. Our results indicate that the assembly of cytoophidium complexes with different compositions resonates with distinct cell fates.

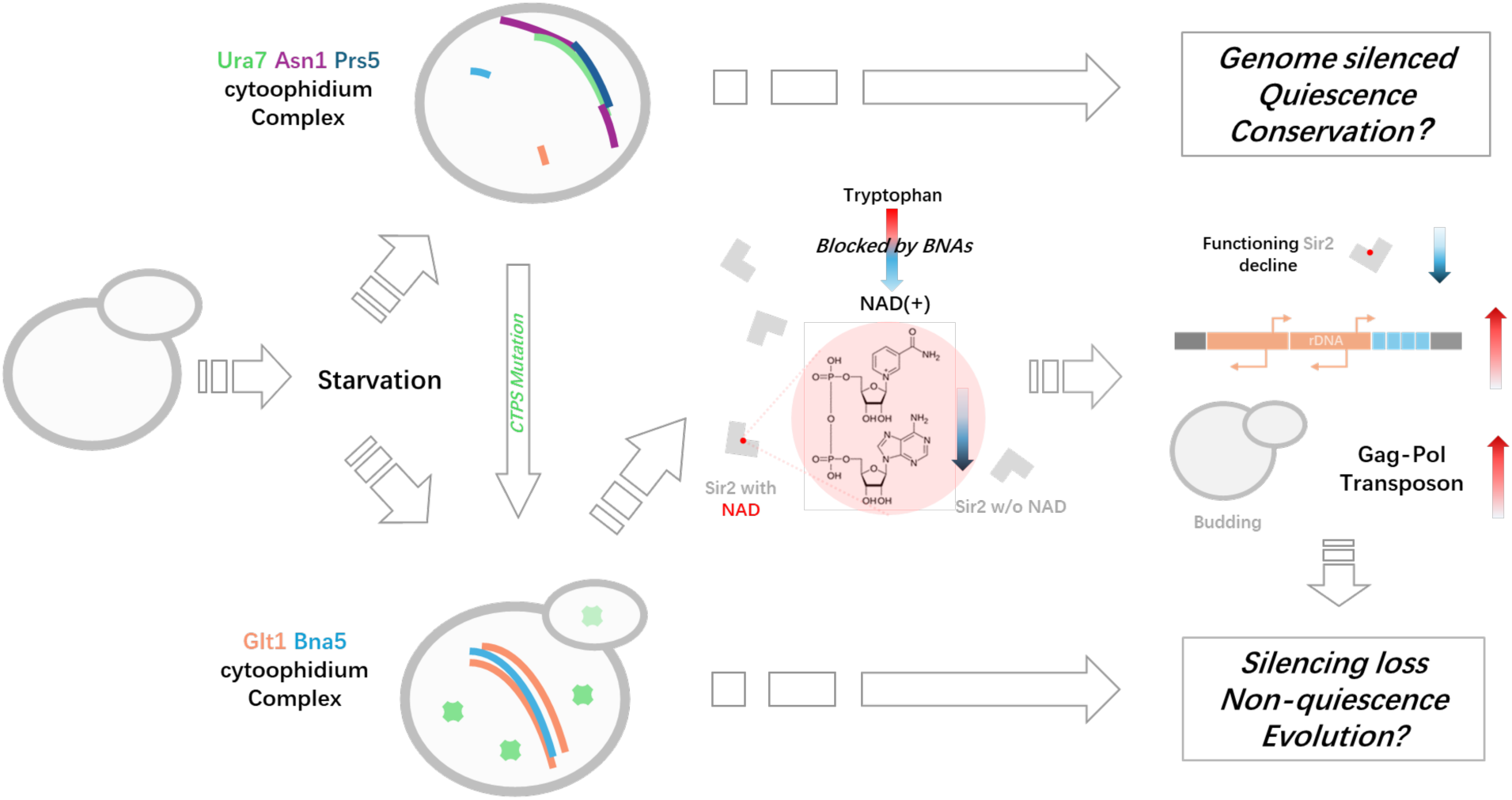

**HIGHLIGHTS:**

- Multiple cytoophidia can be assembled into cytoophidium complexes by docking one after another.
- CTPS cytoophidia blockage impedes quiescence and boosts Glt1-Bna5 cytoophidium complex.
- Glt1 cytoophidium blockage restores quiescence and Bna5 inhibition in NAD pathway.
- Activating NAD utilizing SIR2 rescues CTPS cytoophidium blockage induced non-quiescence.

## INTRODUCTION

Metabolism is one of the most fundamental characteristics of life, determining the cell fate of replication, quiescence, senescence and tumorigenesis[1–7]. Thus, the regulation of metabolic enzymes is critical to fate determinations[8–10]. In 2010, we reported that cytidine 5’-triphosphate synthase (CTPS) forms filamentous structure, termed cytoophidia (cell serpents in Greek) in *Drosophila*[11]. Subsequently, similar phenomenon was reported in bacteria [12], budding yeast and rats [13].

In *Drosophila*, the CTPS cytoophidium enhances ammonia channeling efficiency[14]. However, in budding yeast, cytoophidia act as a storage strategy, reducing activity during starvation[15]. Several non-enzymatic functions have also been identified, including prolonging enzyme half-life[16], regulating cell adhesion[17], and influencing lipid metabolism via the integrin signaling pathway[18]. Additionally, the CTPS cytoophidium is associated with tumorigenesis[19, 20] and thymocyte maturation[21, 22]. The cytoophidium-forming ability of CTPS is highly conserved across all three domains of life[12, 23–27].

Interestingly, CTPS is not the only cytoophidium-forming enzyme. Dozens of such enzymes have been found in localization screenings in *Saccharomyces cerevisiae*[13, 28, 29] and other studies[22]. This underscores the prevalence of cytoophidia as a previously overlooked mode of metabolic enzyme organization. However, the spatial and functional relationship of cytoophdium-forming enzymes remain elusive.

To systematically study cytoophidium-forming enzymes, we take advantage of the model organism *Saccharomyces cerevisiae* for its wealth of optical and genetical tools. In this study, we focus on five metabolic enzymes that form cytoophidia: ASNS (Asn1 in budding yeast), CTPS (Ura7), GOGAT (Glt1), PRPS (Prs5), and kynureninase (BNA5). Super-resolution live-cell imaging allows us to observe cytoophidium formation in real-time. In addition, we introduce mutations that inhibit cytoophidium formation for phenotype analysis, paving the way for high-throughput investigations through transcriptomics, metabolomics and the following verifications.

We reveal that multiple cytoophidia can organize into cytoophidium complexes through sequential docking. Notably, Glt1 cytoophidia predominantly assemble in non-quiescent cells, while CTPS cytoophidia are more prevalent in quiescent cells, forming complexes with Prs5 and Asn1 cytoophidia. Disrupting CTPS cytoophidium assembly can induce a non-quiescent phenotype and promote the assembly of Glt1 and Bna5 cytoophidia and their complexes. This disruption also inhibits the NAD biosynthesis pathway involving Bna5 and the silencing regulator Sir2. Importantly, the non-quiescent phenotype can be rescued by inhibiting Glt1 assembly, supplementing nicotinic acid, or overexpressing Sir2, highlighting that cytoophidium complexes resonate with cell fates.

## RESULTS

### Multiple metabolic enzymes can form cytoophidium complexes

In order to study the interactions of cytoophidia in a systematic manner, four proteins of interest were labeled in a single budding yeast cell, specifically Prs5 mTagBFP (pyrophosphate synthase, PRPS), Ura7 mGFP (cytidine triphosphate synthase, CTPS), Glt1 mCherry (glutamate synthase, GltS), Asn1-miRFP670nano (asparagine synthase, ASNS). This strain was named UGPA. During a 5-day fixed culture, all labeled enzymes were able to form linear structures at the micrometer level (Fig. 1 A-D), indicating cytoophidia. Some cytoophidia can also attach to each other, showing the characteristic of forming complexes (Fig. 1 E). The statistical data of the complex shows that nearly 60% of cells (cells with internal cytoophidia) do not have cytoophidium complexes, while nearly 40% of cells do have cytoophidium complexes (Fig. 1 G, I).

**Figure 1.**
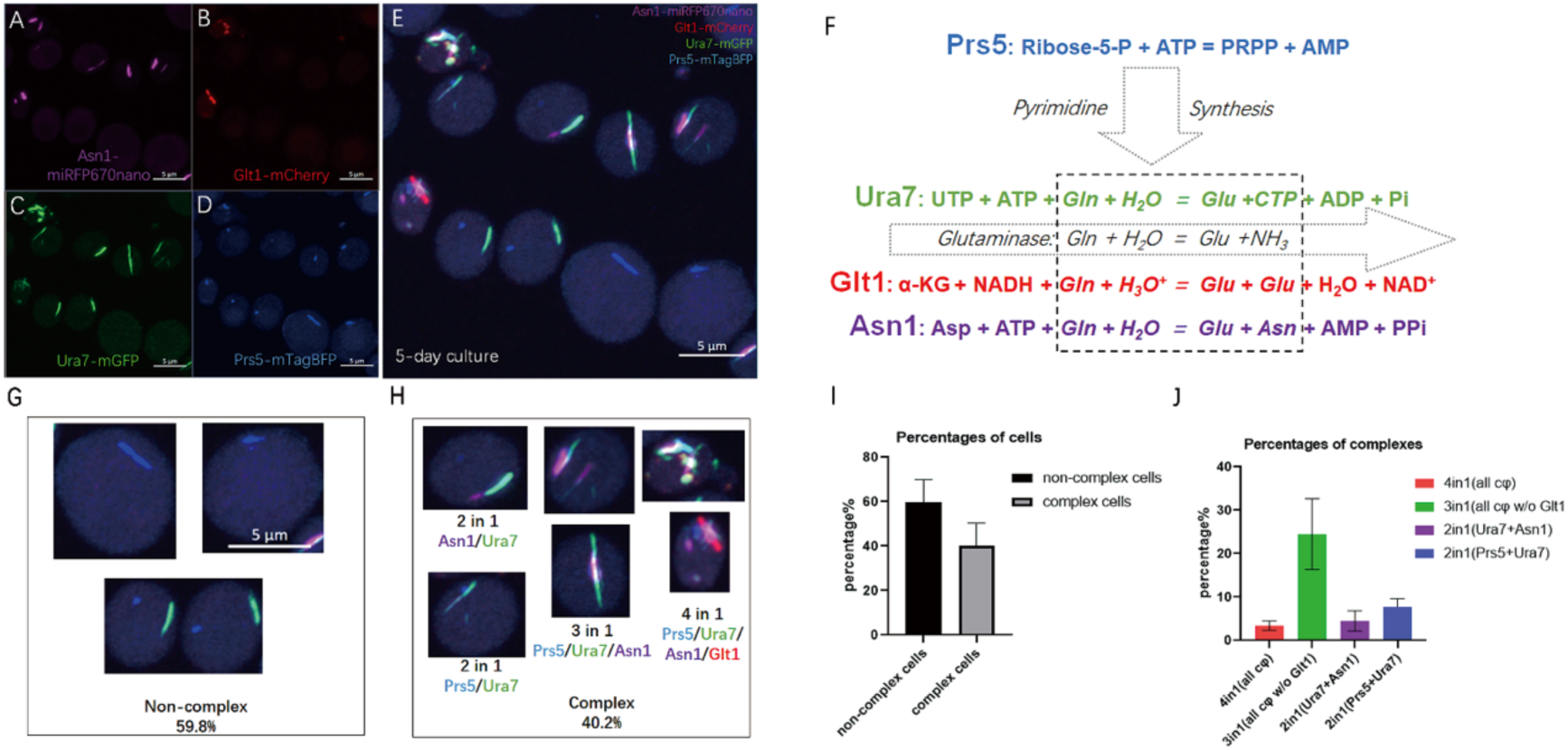
Multiple metabolic enzymes can form cytoophidium complexes. **A-D.** Different channels of representative cytoophidium complexes, purple represents Asn1-miRFP670nano (A), red represents Glt1-mCherry (B), green represents Ura7-mGFP (C), blue represents Prs5-mTagBFP (D). **E.** Overview of cytoophidium complex in 5-day culture. **F.** The reaction of the enzymes of interest. α-KG = α-ketoglutarate, Asp = aspartate, Asn = asparagine, Glu = glutamate, Gln = Glutamine. **G-H.** Illustration of non-complex cytoophidia (G) and different cytoophidium complexes (H). **I-J.** Quantification of the percentage of cells containing or not containing cytoophidium complexes (I) and the percentage of complexes of different compositions (J). Scale bar = 5 μm. Cells in F and G are in the same scale.

According to the types of cytoophidium composed in the complexes, they can be divided into three categories: One complex contains two cytoophidia (2in1), One complex contains three cytoophidia (3in1) and one complex contains four cytoophidia (4in1). It is interesting that the probability of 2in1 and 3in1 cytoophidium combinations is not equal. If it is a random event, there are only 3 types in total, not 10. The preference of the cell finally leads to 4 types of cytoophidium complexes, namely Asn1/Ura7 (2in1), Prs5/Ura7 (2in1), Prs5/Ura7/Asn1 (3in1), and Prs5/Ura7/Asn1/Glt1 (4in1) (Fig. 1 H). In stationary culture, the 3in1 complex is the most abundant, while the 4in1 complex is the least abundant. Interestingly, Ura7 is the only cytoophidium present in all complexes. Combining the bridging position of Ura7 between pyrimidine synthesis and glutamine consumption, Ura7 may play a role in complex formation (Fig. 1 F).

To eliminate the bias induced by fluorescent protein tagging, the combinations of fluorescent proteins and enzymes are switched to Ura7-mTagBFP, Glt1-mGFP, Asn1-mCherry, and Prs5-miRFP670nano. This switched strain is named GAUP, and similar cytoophidium complexes have also been found in GAUP. (Fig. S1).

### Cytoophidia form complex by docking one-by-one

To investigate the formation of cytoophidium complexes over time, cultures were used at different time points in log phase (8 hours), diauxic phase (1 day), and stationary phase (5 day). Data showed that Prs5, Ura7, and Asn1 exhibited a starvation-induced cytoophidium forming pattern, while Glt1 did not and only showed a relatively low abundance, up to about 10% (Fig S2. A, B). This was much lower than previously reported. The reason might be that the dimerization of GFP tagging amplified the filamentation effect of the enzymes, while the monomeric version used in this research did not.

For the complexes, the changes of the abundances are more diverse (Fig. S2 C). The 2in1(Ura7/Prs5) complex decreased from the diauxic phase to the stationary phase, while the abundance of the 3in1 complex increased significantly, indicating a transformation of the 2in1(Ura7/Prs5) complex into the 3in1 complex. However, the abundance of the 2in1(Ura7+Asn1) complex only slight increased, indicating that Asn1 and Ura7 cytoophidia were more inclined to bind to other complexes. Due to the low abundance of Glt1 cytoophidium itself, it was expected that the 4in1 complex would slightly increase over time.

To gain a deeper understanding of the transformation process of the complex, we performed live imaging of strains UGPA and GAUP under starvation induction. Log state active cells were induced from old medium cultured for 5 days and immediately imaged (within 10 minutes). During the imaging process, only Prs5, Ura7, and Asn1 cytoophidia and their related complexes were recorded, but no Glt1 cytoophidium was detected (Fig. 2A). This result confirms that the formation of Glt1 is not starvation dependent.

**Figure 2.**
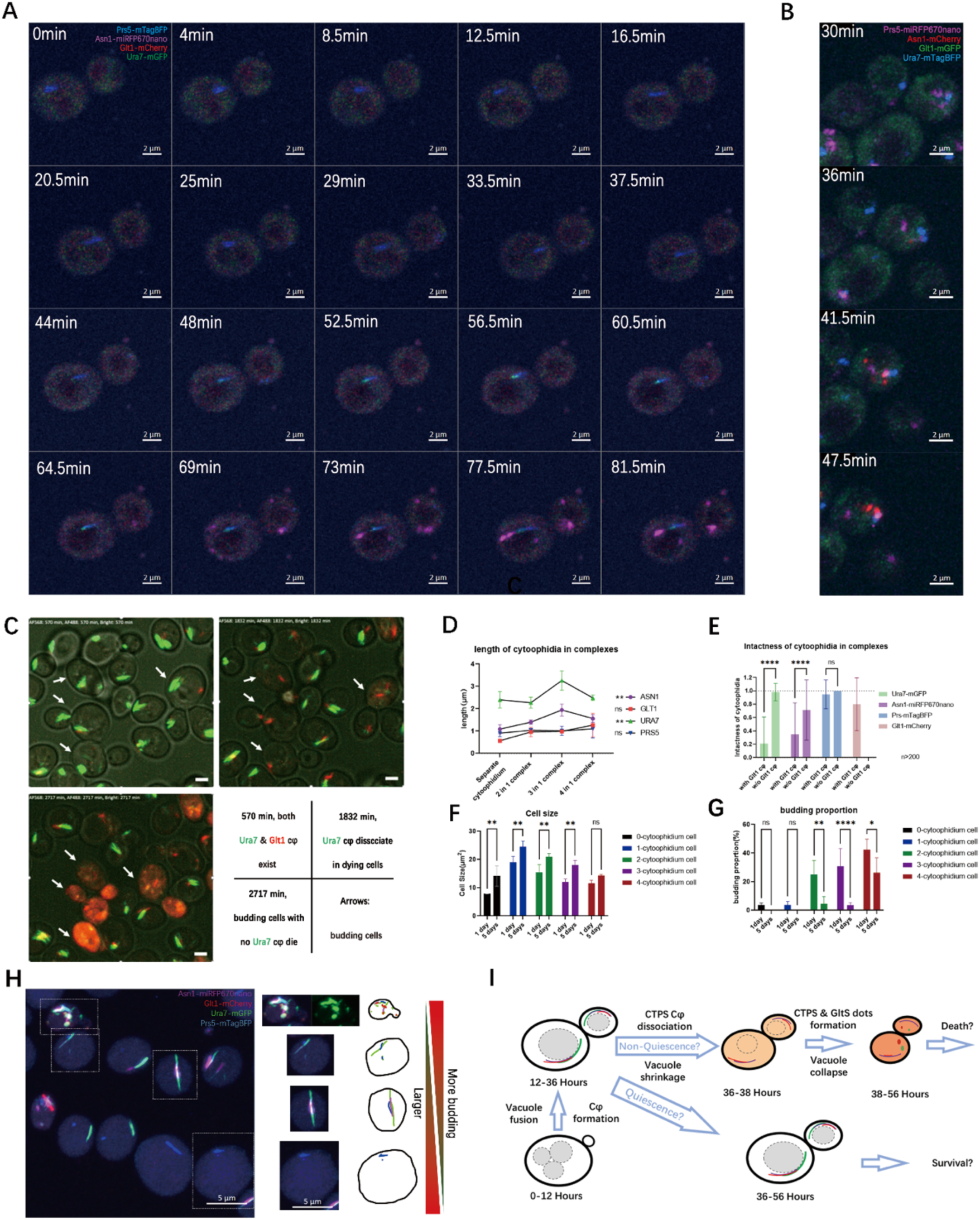
Live imaging of complex formation reveals the non-quiescent phenotype associated with Glt1-Ura7. **A.** Time series of cytoophidium complex formation induced by old culture medium in UGPA strains. The time marked in the upper left corner represents the time since the capture began. Purple represents Asn1-mRFP670nano, red represents Glt1 mCherry, green represents Ura7mGFP, and blue represents Prs5-mTagBFP. Scale bar = 2 μm. **B.** Time series of cytoophidium complex formation induced by old culture medium in GAUP strains. The time marked in the upper left corner represents the time since the capture began. Purple represents Prs5-miRFP670nano, red represents Asn1 mCherry, green represents Glt1 mGFP, and blue represents Ura7 mTagBFP. Scale bar = 2 μm. **C.** The key framework for Glt1 cytoophidium formation, Ura7 cytoophidium dissociation, and cell death. The status and time of keyframes are displayed in the lower right corner of the panel. **D.** Quantification of the length of different cytoophidia in composite complexes. **E.** Quantification of Ura7 cytoophidium integrity in cells with and without Glt1 cytoophidia. **F.** Cells with different complexes have different sizes. 4-cytoophidium cells with Glt1 cytoophidia have the smallest average size. **G.** The budding rate of cells with Glt1 cytoophidia is the highest. Scale bar = 5 μm. **H.** An overview was given of the heterogeneity in length, cytoophidium integrity, cell size, and budding rate of cytoophidium complexes with different components. **I.** A cartoon model showcasing the relationship between cytoophidium interactions and cell fate determination. * indicates p-value < 0.05; **, p-value < 0.01; ***, p-value < 0.001; ****, p-value < 0.0001; and ns or unmarked indicates no significance in the statistical chart.

Live cell imaging also showed that during starvation, cytoophidia formed in the order of Prs5, Ura7, and Asn1. In strain UGPA (Fig. 2A), Prs5-mTagBFP formed aggregates very quickly after induction and before imaging began (within 10 minutes). Then, the elongation of Prs5-mTagBFP cytoophidia lasted for nearly 45 minutes, and Ura7-mGFP aggregates appeared, docking with the end of Prs5 cytoophidia. The elongation lasted for 20 minutes, forming a partially colocalized complex of Ura7 and Prs5. Around 69 minutes after the start of imaging, Asn1-miRFP670nano aggregated into multiple foci. One of the Asn1 foci docked at one end of the complex, and eventually more foci aggregates there. However, during the 180-minute induction process, no Glt1 cytoophidium was found. The live imaging of strain GAUP also showed similar results (Fig. 2 B).

Live imaging revealed that cytoophidia formed in the order of Prs5, Ura7 and Asn1 under starvation, transitioning from a 2in1(Prs5/Ura7) to a 3in1 complex in a dock-and-grow manner. However, Glt1 cytoophidium could not be induced through simple starvation induction, implying the specialty and differences in the interaction between other cytoophidia and the Glt1 cytoophidium.

### Glt1 cytoophidium suppresses Ura7 cytoophidium and highlights non quiescent cells

To reveal how Glt1 cytoophidium is formed, we used 3-day live cell imaging in YPD (Fig. 2C). Live imaging showed that Glt1 cytoophidia could form during the standard culture process after Asn1 cytoophidium formation (Fig. 2C top left and Fig. S3A). However, over time, Ura7 cytoophidia dissociated in the budding cells with Glt1 cytoophidia (Fig. 2C top right and Fig. S3B). Finally, the budding cells with dissociated Ura7 cytoophidia died (Fig. 2C bottom left and Fig. S3B). This indicates that the formation of Glt1 cytoophidium is opposite to that of Ura7 cytoophidium.

In addition, live imaging captured the process of Asn1 foci fusing into elongated and larger structures, forming mature Asn1 cytoophidia at the location of Ura7 cytoophidia (Fig. S3A). It was worth noting that after Ura7 cytoophidia dissociation, Glt1 cytoophidia and Asn1 cytoophidia still maintained contact. Although Ura7, Glt1, Asn1 aggregated again into a large complex after cell contraction and death (Fig. S3B), this transient Glt1-Asn1 cytoophidium complex indicated a closer relationship between Glt1 and Asn1 cytoophidia. Due to the low sensitivity of the camera, Prs5 labeled with blue fluorescent protein was almost unrecognizable in long-term live imaging.

Due to the significant differences between Glt1 cytoophidium and other cytoophidia, we expected cells containing it to have more special characteristics (4in1 complex cells). We quantified the lengths of Asn1, Glt1, Ura7 and Prs5 cytoophidia in various complexes and isolated cytoophidia (Fig. 2D). For Glt1 and Prs5 cytoophidia, the length was always around 1 micrometer. However, for Asn1 and Ura7 cytoophidia, when Glt1 cytoophidia involved in the 4in1 complex, the length decreased, indicating that Glt1 cytoophidium hindered the formation of Asn1 and Ura7 cytoophidia.

To investigate this possible inhibitory effect, cells with 4in1 complex were examined. Dispersed and shortened Ura7 cytoophidia were found in the 4in1 complex (Fig. 2H), rather than individual intact cytoophidia in other cells. The integrity of cytoophidia (1 is greater than the number of cytoophidium in a cell) indicated that compared to cells without Glt1 cytoophidia, Ura7 and Asn1 cytoophidia were losing integrity in cells with Glt1 cytoophidia. The integrity of Prs5 cytoophidium was not affected by the presence of Glt1 cytoophidium (Fig. 2E). It seemed that Glt1 cytoophidium was related to the dissociation of Ura7 and Asn1 cytoophidia, but not that of Prs5 cytoophidium.

The cell morphology also varied among cells with different cytoophidium combination, showing a decreasing size when a cell has more types of cytoophidia (Fig. 2F). When comparing the sizes of 1-day and 5-day cells, most cells grew larger except for the smallest 4-cytoophidium cells. This suggests that 4-cytoophidium cells containing Glt1 cytoophidium are possibly dividing cells, but not swollen G0 quiescent cells. In 1-day of culture, the budding rate of 4-cytoophdium cells (Cells containing Glt1 cytoophidium) was quite high, about 40% (Fig. 2G). Although 2-cytoophidium cells and 3-cytoophidium cells had a budding rate of nearly 30% in 1-day culture, they sharply decreased to less than 5% in 5-day stationary culture. However, for 4-cytoophidium cells, the budding rate was still high, at 25%, showing non-quiescent characteristics.

To elucidate this complex relationship, we established a cartoon model for cytoophidium-coupled cell fate transition (Fig. 2I), indicating the connection between cytoophidium combinations and possible cell quiescence determination.

### CTPS cytoophidium disruption leads to more Glt1 cytoophidia and impairs cell quiescence

Due to the correlation between CTPS cytoophidium integrity and cell fate, we planned to manipulate cytoophidia and quiescence entry to verify their correlation. Considering the difficulty of triggering more Glt1 cytoophidium alone and the relatively easy method of manipulating CTPS cytoophidium formation with the help of structural research of Ura8, we have developed a mutant strain URA7(H360A, D370A, R391A, W392G)-mGFP, URA8(H360A, D370A, K391A, W392A)-mCherry (CTPS mutant strain), which cannot aggregate CTPS into cytoophidia. On the other hand, acetyltransferase ARD1 deletion can lead to G0 entry failure, making it a good method to verify the effect of quiescence on CTPS cytoophidium.

The results showed that, as a non-quiescent control strain, ARD1 deletion cells had a higher budding rate and lower Ura7 cytoophidium abundance compared to the control strain in 1-day culture, while the CTPS mutant had a higher budding rate and no CTPS cytoophidia (Fig. 3A, C). In 5-day saturated culture, the ARD1 deletion strain showed no significant difference in Ura7 cytoophidium abundance compared to the control group, but still had high budding rate and Glt1 cytoophidium abundance. The CTPS mutant showed a high budding rate and Glt1 cytoophidium abundance similar to the ARD1 deletion strain (Fig. 3D), indicating that CTPS cytoophidium disrupting mutations have a strong fate-determining effect of the. By the way, the enhanced Glt1 cytoophidium in CTPS mutants and ARD1 deletion strains can also be considered as non-quiescence related markers.

**Figure 3.**
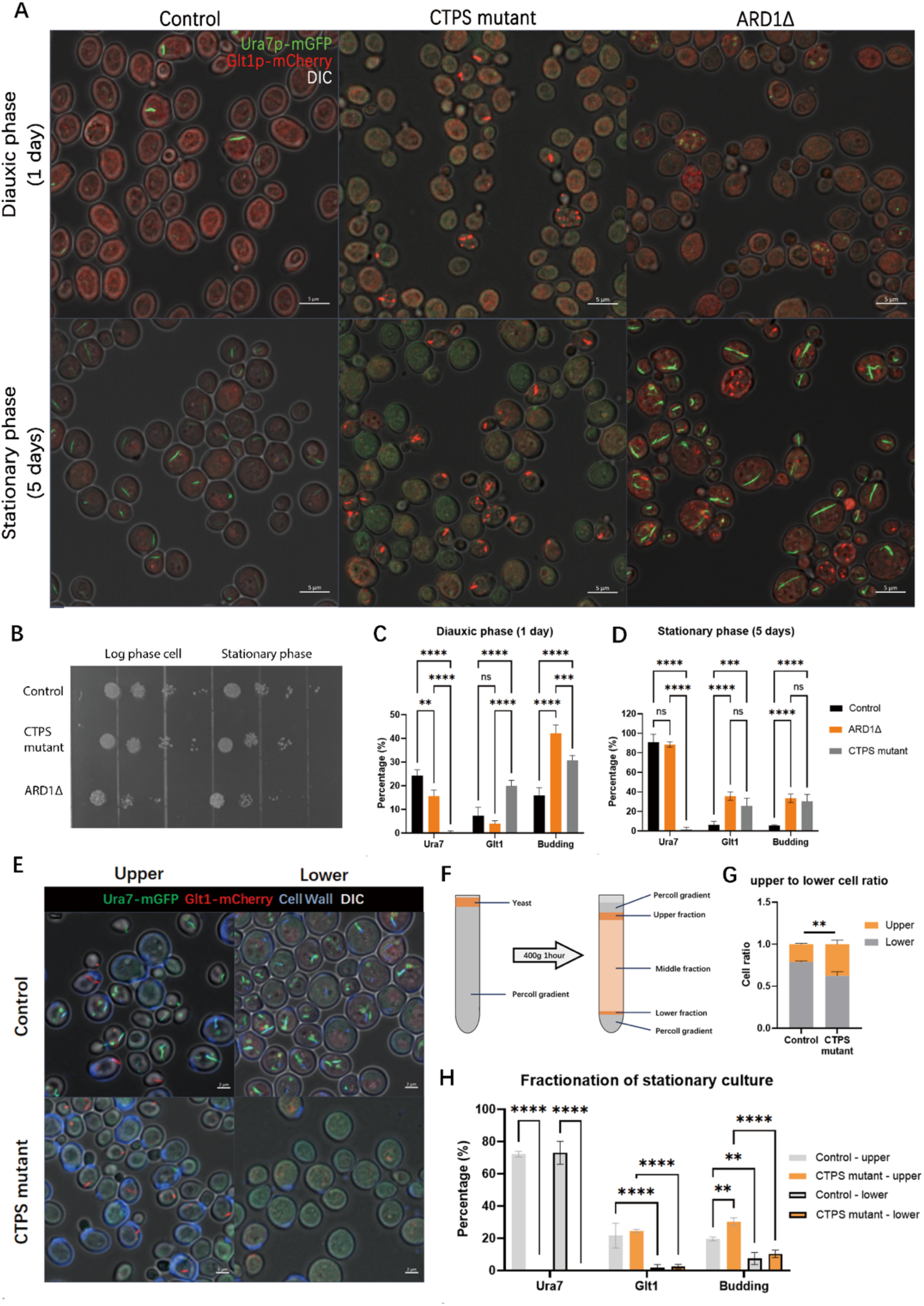
CTPS cytoophidium disruption leads to more Glt1 cytoophidia and impairs cell quiescence. **A.** Confocal images of control strain (UGPA), CTPS mutant (Ura7mutant-mGFP, Ura8mutant-miRFP670nano, Glt1-mCherry) and ARD1Δ (Ura7-mGFP, Glt1-mCherry, ARD1Δ) during diauxic phase (1 day) and stationary phase (5 days). Only Ura7 signal (green), Glt1 signal (red) and DIC (white) are shown. Scale bar = 5 μm. **B.** Spot assay of strains in A, spotted at log phase and stationary phase. **C-D.** Percentages of Ura7 cytoophidium and Glt1 cytoophidium positive cells and budding rate of the strains mentioned in A, at diauxic phase (1 day, C) and stationary phase (5 days, D). **E.** Confocal images of the fractionated control strain and CTPS mutant mentioned in A. Only Ura7 signal (green), Glt1 signal (red), CFW signal (cell wall, blue) and DIC (white) are shown. Scale bar = 2 μm. **F.** Procedure of Percoll based cell fractionation. **G.** Cell ratio of different fractions from control strain and CTPS mutant. **H.** Percentages of Ura7 cytoophidium and Glt1 cytoophidium positive cells and budding rate of the strains mentioned in E. * means p-value < 0.05; **, p-value < 0.0; ***, p-value < 0.001; ****, p-value < 0.0001; and ns or not labelled mean no significance in statistical chart.

Nonetheless, CTPS mutants also exhibited several other non-quiescent phenotypes, including low survival rate when inoculate at stationary phase (Fig. 3B), higher ROS level (Fig. S4A, B), more older cells (Fig. S4C) and shorter lifespan (Fig. 3D). In many previous publications, all these phenotypes were typical manifestations of non-quiescent cells, strongly indicate that CTPS cytoophidium disrupting mutations can lead to non-quiescence fate.

To distinguish the effects of cytoophidium formation, expression levels, and enzyme activity on non-quiescent phenotypes, CTPS overexpression strains were cultured, and 1 mM cytosine was added to control strains (Fig. S5). The results showed that overexpression of the CTPS mutant still exhibited the same high budding rate and Glt1 cytoophidium abundance, with no significant difference compared to the CTPS mutant knock-in strain (Fig. S5 A-C.). After adding 1 mM cytosine to the control strain, there was no difference in Glt1 cytoophidium abundance and budding rate compared to the control strain (Fig. S5 A-C.). These results indicate that CTPS expression levels and additional cytosine nucleotide base are not determining factors for the non-quiescent phenotypes of CTPS mutants.

In order to gain a deeper understanding of the details between CTPS mutants and non-quiescence, Percoll gradient based fractionations were performed on the stationary phase control and CTPS mutants. The density of quiescent cells is higher and will go lower, while non-quiescent cells are lighter and will remain upper during the fractionation process (Fig. 3F). The results showed that the Glt1 cytoophidium abundance and budding rate in upper fractions (non-quiescent cells) were higher than those in the lower fraction (quiescent cells) (Fig. 3E, H). The similarity between the upper fractions of these two strains once again indicates that Glt1 is a marker of highly budding, lighter, non-quiescent cells. Afterwards, the difference of quiescence between the control strain and the CTPS mutant was attributed to the ratio of upper and lower cells. The quantification of cell ratio showed that the CTPS mutant had more non-quiescent cells than the control strain, resulting in a significant non-quiescent phenotype (Fig. 3G).

### CTPS cytoophidium disruption blocks NAD synthesis and **impedes cell silencing**

To understand the mechanism by which CTPS mutants lead to non-quiescence, transcriptome studies combined with metabolome were conducted. The log and stationary phases of the control and CTPS mutant strains were used for transcriptome analysis, and PCA clustering was used to display different cell fates starting from similar origins (Fig. 4A). The metabolomes of the two strains of stationary phase were also fall into two separate clusters (Fig. 4B).

**Figure 4.**
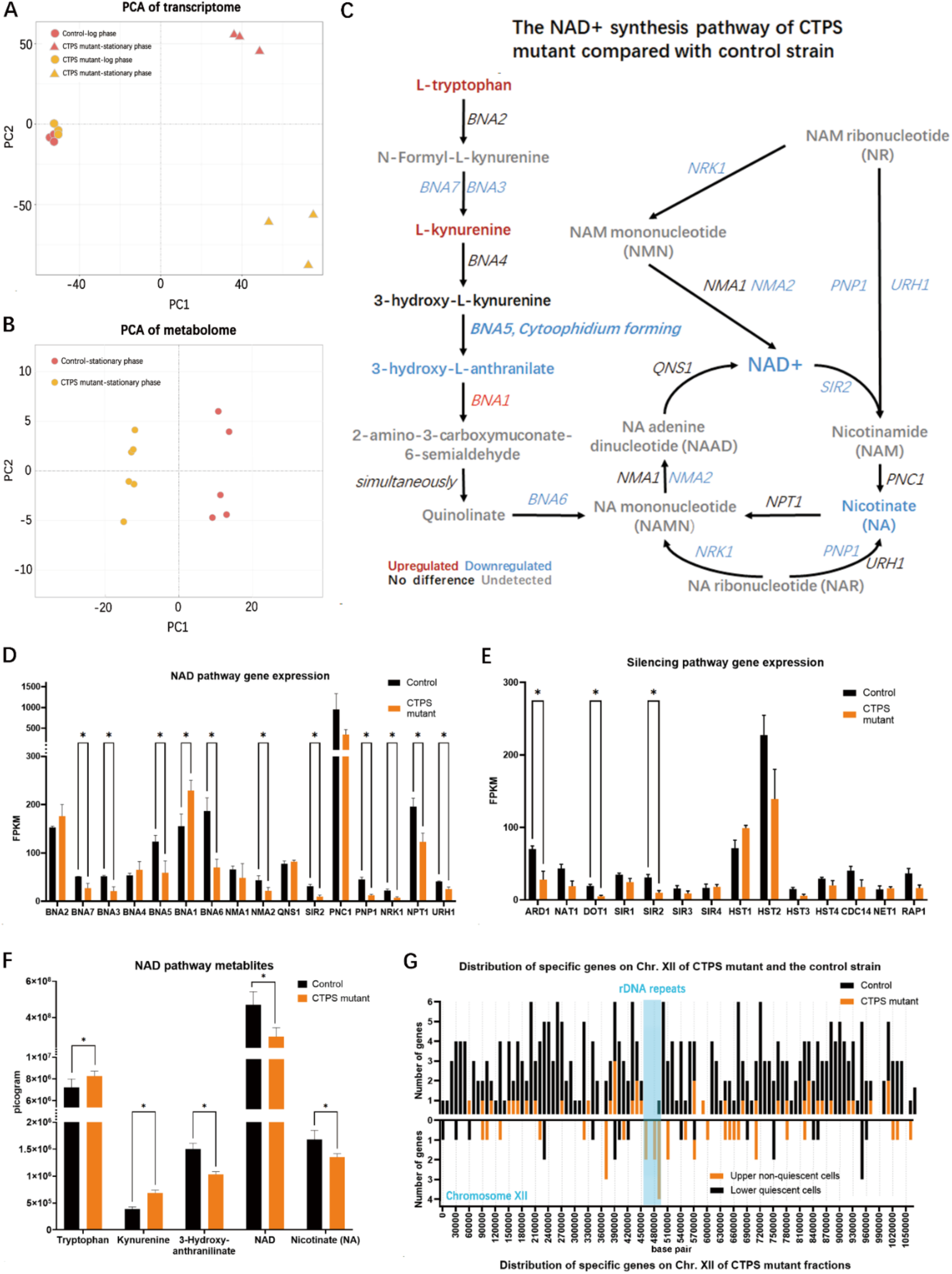
CTPS cytoophidium disruption blocks NAD synthesis and impedes cell silencing. **A.** Transcriptome PCA plots of UGPA and CTPS mutant (Ura7^mutant^-mGFP, Ura8^mutant^-miRFP670nano, Glt1-mCherry) in log phase (8 hours) and stationary phase (5 days). **B.** Metabolome PCA plots of UGPA and CTPS mutant mentioned in A are only in the stationary phase. **C.** NAD (+) synthesis pathway, labelled with differential expressed genes and metabolites, from the comparisons of stationary phase multi-omics between control and CTPS mutant. **D.** Comparison of expression levels of NAD synthesis pathway genes between control and CTPS mutant in the stationary phase. **E.** Comparison of expression levels of silencing related genes between control and CTPS mutant in the stationary phase. **F.** Comparisons of absolute amount of metabolites in NAD synthesis pathway between control and CTPS mutant in the stationary phase. **G.** The distribution of differential expressed genes on Chromosome XII shows the differentially expressed genes between control and CTPS mutant (upper part) and differentially expressed genes between upper and lower cells of CTPS mutant (lower part) in the stationary phase. Blue square represents the location of rDNA repeats. * means p-value < 0.05; **, p-value < 0.01; ***, p-value < 0.001; ****, p-value < 0.0001; and ns or not labelled mean no significance in the statistical chart.

After comparing the two omics combinations, it was found that the NAD superpathway is one of the common differences in transcription and metabolism (Fig. 4C, D, F). The NAD superpathway consists of the NAD *de novo* synthesis pathway and the salvage pathway. For the de novo pathway, the BNA (Biosynthesis of nicotinic acid) genes are responsible for converting the amino acid tryptophan into nicotinate mononucleotide (NAMN), which is a precursor to NAD. It was very interesting that the products of BNA first surged in kynurenine and then decreased in 3-hydroxy-L-anthranilate (Fig. 4C, F). The gene responsible for this transition is kynureninase BNA5, which, like most other BNA genes, only showed half level gene expression in transcription (Fig. 4D). This clue suggests that BNA5 acts as a valve in de novo NAD synthesis. Moreover, BNA5 can even form cytoophidia, indicating a connection between metabolic valving and the cytoophidium.

For the NAD salvage pathway (genes other than BNAs in Fig.4C), most genes were down regulated, including the NAD(+)-dependent silencing information regulator SIR2, which is a key deacetylase for genome silencing and longevity (Fig. 4C, E). In addition to SIR2, some other silencing related genes in CTPS mutants were down regulated, including ARD1 and DOT1 (Fig. 4E). ARD1 has been introduced in previous results, which is a key regulator for cell quiescence. The downregulation of ARD1 in CTPS mutants corresponds to the deletion of ARD1, which may partially explain the non-quiescent phenotype of CTPS mutants. Moreover, the acetyltransferase activity of ARD1 is also crucial in SIR2 silencing complex targeting chromosomes[30]. Interestingly, this targeting mechanism requires another factor, histone methylase DOT1[30, 31], which was also down regulated in CTPS mutants (Fig. 4E). In summary, the downregulation of three silencing regulatory factors, SIR2, ARD1, and DOT1, in CTPS mutants indicates the presence of genomic silencing defects.

To confirm the possibility of silencing loss, we mapped the positions of differentially expressed genes in the transcriptome data onto chromosome XII, where rDNA repeat sequences are located (Fig. 4G). The mapping showed that when comparing the control strain and CTPS mutant, only one gene is differentially expressed in the rDNA region, indicating a normalized result. However, when comparing the upper and lower fraction of the CTPS mutant, 8 genes were specifically expressed in the upper fraction cells of the rDNA regions compared to the lower quiescent fraction. The 8 upregulated rDNA genes were TAR1, YLR154C-G, YLR157C-B, YLR157W-E, YLR159C-A, ASP3-4, YLR162W and RRT15. Besides, the Gag-Pol transposons also get upregulated in the upper fraction of the CTPS mutant (Fig. 6D), which may contribute to evolution[32–34].

In summary, by combining transcriptome and metabolome analysis, it was found that the CTPS mutant exhibited defects in the shut-down of the NAD superpathway and rDNA silencing. BNA5, the cytoophidium-forming kynureninase, can be responsible for the shut-down of NAD synthesis due to its controlled metabolite flux narrowing. Meanwhile, the downregulated NAD-dependent silencing regulator SIR2 may be the ultimate effector downstream of BNAs, leading to a non-quiescent state.

### Glt1 cytoophidium disruption alleviates non-quiescence, related **to NAD synthesis**

Glt1 cytoophidium is highly abundant in CTPS mutants as well as in the upper non-quiescent cells, indicating that it is a hallmark of non-quiescent cells. Therefore, disrupting Glt1 cytoophidium is a good attempt to rescue non-quiescent phenotypes.

To verify the role of Glt1 cytoophidium in quiescence, we developed a Glt1 mutant strain with CTPS mutant background. This mutant showed a significant decrease in Glt1 cytoophidium abundance, which has a good rescue effect on highly budding phenotypes, and a higher proportion of elderly cells in diauxic and stationary phases of cells (Fig. 5A-C). To investigate the following mechanism, we examined published transcriptome data and compared the stationary Glt1 mutant and wild type[35]. Surprisingly, the Glt1 mutant showed significant upregulation of NAD synthesis pathway genes, including most BNA genes and remained salvage pathway expression (Fig. 5D, F). Since the CTPS mutant showed downregulation in the salvage pathway compared to the wild-type, the Glt1 mutant should restore the downregulated salvage pathway to the same level as the wild-type.

**Figure 5.**
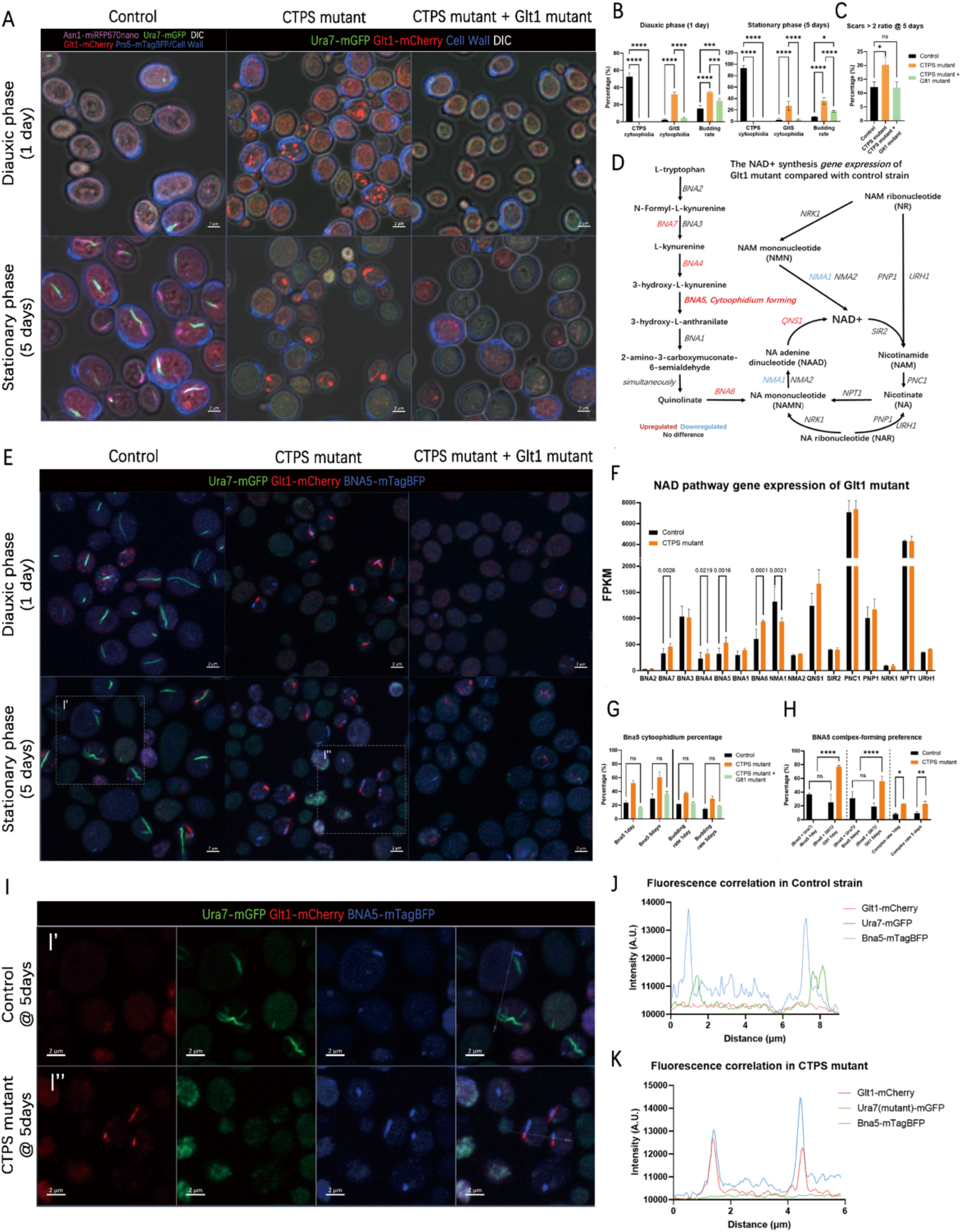
Glt1 cytoophidium disruption alleviates non-quiescence through NAD synthesis related mechanisms. **A.** Confocal images of control strain (UGPA), CTPS mutant (Ura7^mutant^-mGFP, Ura8^mutant^-miRFP670nano, Glt1-mCherry) and CTPS mutant + Glt1 mutant (Ura7^mutant^-mGFP, Ura8^mutant^-miRFP670nano, Glt1^mutant^-mCherry) at diauxic phase (1 day) and stationary phase (5 days). For the control strain with complex, Ura7 signal (green), Glt1 signal (red), Prs5-mTagBFP /CFW (blue), Asn1-miRFP670nano and DIC (white) are shown. For the mutants, only Ura7 signal (green), Glt1 signal (red), CFW (cell wall, blue) and DIC (white) are shown. Scale bar = 2 μm. **B.** The percentages and budding rate of Ura7 cytoophidium and Glt1 cytoophidium positive cells in diauxic phase (1 day, left) and stationary phase (5 days, right) of the strains mentioned in A. **C.** Ratio of cells with 2 or more scars at 5 days in the strains mention in B. One-way ANOVA is performed for significance analysis. **D.** Stationary phase transcriptome comparison between control and Glt1 mutant from open resource using NAD (+) synthesis pathway marked with differentially expressed genes. **E.** Confocal images of control strain (Ura7-mGFP, GlT1-mCherry, Bna5-mTagBFP), CTPS mutant (Ura7^mutant^-mGFP, Ura8^mutant^-miRFP670nano, Glt1-mCherry, Bna5-mTagBFP) and CTPS mutant + Glt1 mutant (Ura7^mutant^-mGFP, Ura8^mutant^-miRFP670nano, Glt1^mutant^-mCherry, Bna5-mTagBFP) at diauxic phase (1 day) and stationary phase (5 days). For the control strain with complex, Ura7 signal (green), Glt1 signal (red), Bna5-mTagBFP (blue), and DIC (white) are shown. Scale bar = 2 μm. **F.** Expression level comparison of NAD synthesis pathway genes between control and CTPS mutant at stationary phase. **G,H.** The percentage of Ura7, Glt1, Bna5 cytoophidia positive cells and the percentage of Bna5 cytoophidia in different complexes. **I.** The magnification of the square of the area in E. I’ shows the zoom-in of control strain. I’’ shows the zoom-in of CTPS mutant. Arrows shows the line used for fluorescence correlation analysis in J and K. Scale bar = 2 μm. **J, K.** Fluorescence correlation analysis of lines selected in I’ and I’’, indicating the correlation of cytoophidia in control strain (J) and CTPS mutant (K). * means p-value < 0.05; **, p-value < 0.01; ***, p-value < 0.001; ****, p-value < 0.0001; and ns or not labelled mean no significance in the statistical chart.

It was worth noting that the only cytoophidium-forming protein Bna5 was also upregulated in the Glt1 mutant (Fig. 5F), making it a candidate protein for bridging Glt1 and NAD synthesis. To confirm the effect of Glt1 mutant on Bna5 cytoophidium formation, we developed fluorescent protein tagged control strain, CTPS mutant, and CTPS mutant + Glt1 mutant. Surprisingly, Bna5 cytoophidium showed significant differences among these strains. The CTPS mutant increased the percentage of Bna5 cytoophidium, while the Glt1 mutant completely rescued this increase, as well as the budding phenotype (Fig. 5E, G), indicating the countering regulation of CTPS and Glt1 cytoophidia on BNA cytoophidium.

In addition, in the CTPS mutant, the physical proximity between BNA5 cytoophidium and CTPS or Glt1 cytoophidium also changed (Fig. 5H-K). We found that in the control strain, BNA5 cytoophidium showed no significant preference for forming complexes with Ura7 or Glt1 cytoophidium. However, in the CTPS mutant, Bna5 cytoophidium had no choice but to form a complex with Glt1 cytoophidium at a much higher percentage than the control (Fig, 5H). The zoom-in images of 5-day cells (Fig. 5I) demonstrated the adjacency relationship of Bna5 cytoophidium and other cytoophidia in the control group (Fig. 5I’) and CTPS mutant (Fig. 5I’’). Correlation profiling revealed that in the CTPS mutant, Bna5 signal and with Glt1 signals frequently colocalized, while in the control strain, Bna5 had weak adjacency with Ura7 (Fig. 5 J-K).

To verify the potential determining role of BNA5 in quiescence regulation, a BNA5 knockout strain was also constructed. The results showed that the budding rate of BNA5 knockout strain slightly increased compared to the control at 5 days (Fig. S6 A-C). The partial recurrence of this phenotype indicates that BNA5 is not a direct effector, but rather the presence of another downstream determinant.

Overall, the disruption of Glt1 cytoophidium formation rescued the non-quiescent phenotype induced by CTPS mutants. This rescue effect also provided expression of genes involved in the NAD synthesis pathway, including the key cytoophidium-forming protein Bna5. Bna5 cytoophidium also responded to the CTPS mutant and Glt1 mutant through competitive complex formation. The Bna5 cytoophidium formation deficient strain also exhibited a severe non-quiescent phenotype, indicating that Bna5 plays a role in quiescence regulating along with CTPS and Glt1.

### NAD precursors and NAD-dependent deacetylase SIR2 rescue non-quiescence phenotype

Due to the blockade of the NAD pathway in non-quiescent cells and upregulation in rescued cells, NAD and SIR2, consumers of NAD may play a direct role in quiescent regulation. To verify the role of NAD and SIR2 in quiescent regulation, different precursors and additional copies of SIR2 expression were added to the CTPS mutant.

**Figure 6.**
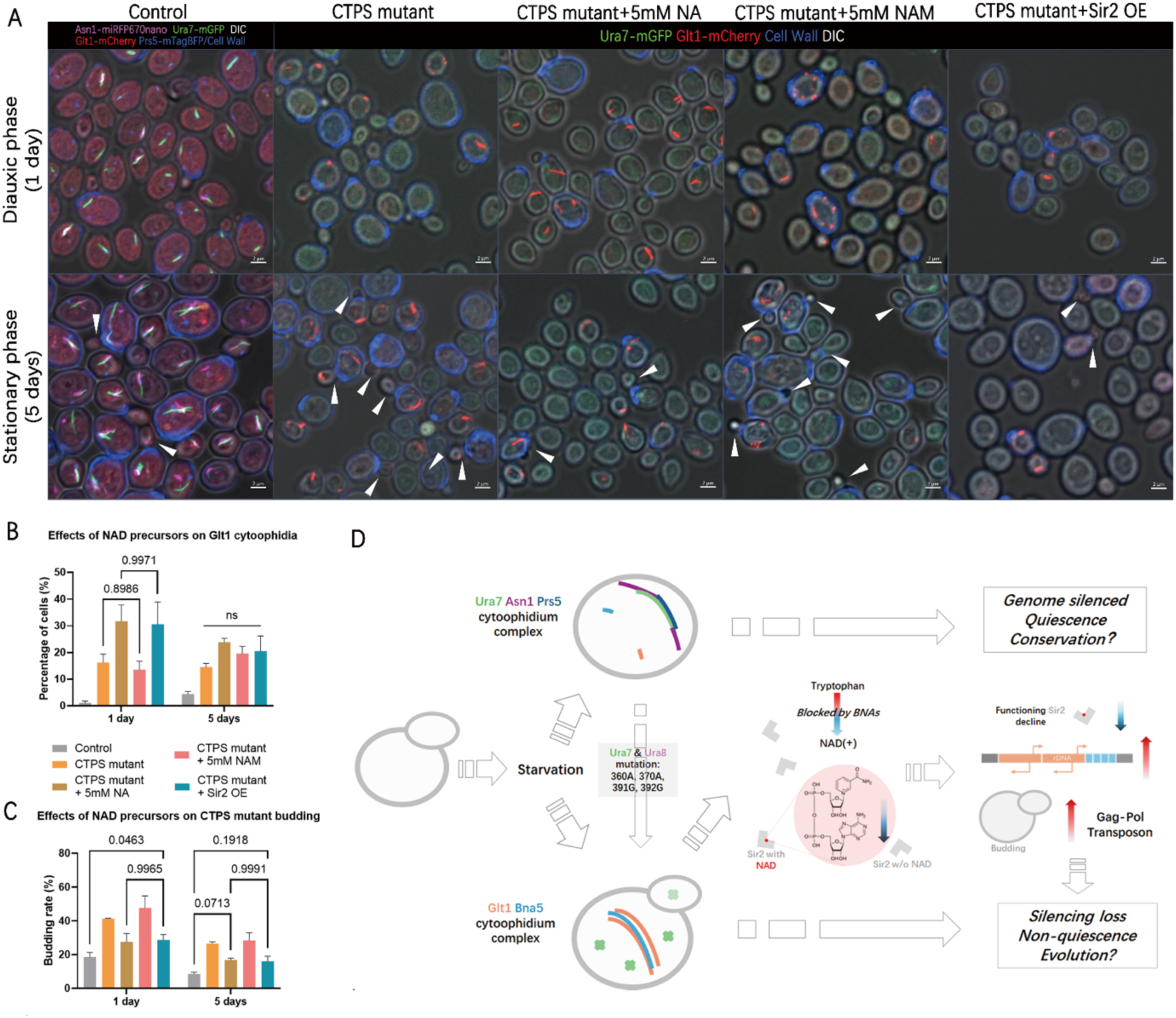
NAD-dependent silencing information regulator SIR2 rescues non-quiescence phenotype. **A.** Confocal images of control strain (UGPA), CTPS mutant (Ura7^mutant^-mGFP, Ura8^mutant^-miRFP670nano, Glt1-mCherry), CTPS mutant with 5mM nicotinate (NA), CTPS mutant with 5mM nicotinamide (NAM) and CTPS mutant + Sir2 OE (Sir2p overexpressed in CTPS mutant strain) at diauxic phase (1 day) and stationary phase (5 days). For the control strain with complex, Ura7 signal (green), Glt1 signal (red), Prs5-mTagBFP /CFW (blue), Asn1-miRFP670nano and DIC (white) are shown. For the mutants, only Ura7 signal (green), Glt1 signal (red), CFW (cell wall, blue) and DIC (white) are shown. Scale bar = 2 μm. Arrowhead indicates buds. **B,C.** The percentage of Glt1 cytoophidium positive cells (B) and budding rate (C) of strains mentioned in A at diauxic phase (1 day) and stationary phase (5 days). **D.** A working model of how cytoophidium interactions lead to different cell fates. p values are indicated as numbers, showing the degree of insignificance.

Two NAD precursors were used in this rescue experiments, both of which can be converted to NAD in the salvage pathway. One is nicotinate (NA, Vitamin B3), and the other is nicotinamide (NAM). The difference between NA and NAM is that NAM is a well-known endogenous SIR2 inhibitor[36, 37] that can help distinguish the effects of NAD levels and SIR2 function. The final concentrations of NA and NAM in YPD were both 5 mM. However, only 5 mM NA showed a rescue effect on the highly budding phenotype of the CTPS mutant, while 5 mM NAM did not (Fig. 6A, C). Furthermore, overexpression of SIR2 in the CTPS mutant also rescued the phenotype, similar to the effect of adding 5 mM NA in diauxic phase (1 day) and stationary phase (5 day) cultures (Fig. 6A, C). This means that not only NAD is critical, but also the functional SIR2 is a determining factor in non-quiescence rescue.

Interestingly, these rescuing manipulations, including NA addition and SIR2 overexpression, promoted the formation of Glt1 cytoophidium during diauxic phase (1 day), while the addition of NAM did not show this effect (Fig. 6B). Considering that Glt1 is also an NAD utilizing enzyme, not only the formation of Glt1 cytoophidium is opposite to that of CTPS cytoophidium, but also SIR2 functions in a competitive manner with NAD. However, during the stationary phase (5 days), this effect became less apparent. In summary, we have created a flowchart model to integrate the results and ideas (Fig. 6D).

## DISCUSSION

This study demonstrates that during starvation, multiple metabolic enzymes can form cytoophidium complexes in a sequential docking manner. We find that there are two main complexes that can be associated with cell quiescence by affecting the biosynthesis of NAD (BNA) (CTPS/ASNS/PRPS complex) or unrelated (GOGAT/Kynureninase complex). Our results reveal the dynamic organization of metabolic enzymes involved in cell fate determination.

### Cytoophidium complexes

Previous studies have shown that different cytoophidia can have pairwise interactions[13, 22, 28, 38, 39]. However, it is unclear whether these pairwise interactions are part of a larger network. In this study, we observe that cytoophidia can form larger complexes and resonate with cell fates. Asn1, Ura7 and Prs5 cytoophidia can form a complex together indicating quiescence, while Glt1 and Bna5 cytoophidia can form a complex indicating non-quiescence. Through live cell imaging, we also reveal that the complex forms in a sequential docking pattern in quiescent cells, with Prs5, Ura7, Asn1 in sequence. However, non-quiescent cells will continue this order with a Glt1 cytoophidium and then dissociate the cytoophidia already formed.

Although our research mainly focuses on the cytoophidium complexes during starvation and subsequent quiescence phenomena, cytoophidia can occur during active growth[28, 40]. For example, cytoophidia discovered in glycolysis pathway[28, 40] can be studied for the presence of their complexes, leading to the development of new functions. The order, preference and circumstances of interactions between different cytoophidia tell us that they are complicated, diverse, and have possible biological functions, rather than just random or physical phenomena. Considering the discovery of a large number of cytoophidium forming proteins in species[29, 41–44], there is great potential for a comprehensive understanding of cytoophidium complexes.

### Determine Cell fate through metabolic means

As is well known, metabolic enzymes can have a determining impact on cell fate, but always in a biochemical manner [2, 3, 10, 45–47]. Here, we propose a cellular biology explanation of how enzymes are physically and visibly organized in determining cell fate, providing a new perspective on the moonlight function of enzymes rather than catalyzing their own reactions. We find that the representative cytoophidia of the two complexes, Ura7 and Glt1cyoophidia, can counteract each other and lead cells towards different fates. CTPS mutant cells with Glt1 cytoophidia exhibit significant classical non-quiescent phenotypes, such as high budding index, high ROS level, low viability, and short lifespan[48–50].

Through multi-omics, we elucidate the critical role of CTPS-GOGAT cytoophidium interactions in determining quiescence in a manner related to the NAD pathway. We find that the biosynthesis of NAD remains at the step of kynureninase Bna5. Bna5 can form cytoophidia and complex with Glt1 cytoophidium in CTPS mutant, while things go the opposite in Glt1 mutant. We speculate that Ura7 and Glt1 cytoophidia control the formation of Bna5 cytoophidia, NAD production and NAD(+) dependent Sir2 activity.

Sir2, also known as sirtuin in mammalian cells, is a key genome silencing regulator, that has been extensively demonstrated to be a controller of cell fate and lifespan[51–53]. Sir2 is an NAD(+) dependent deacetyltransferase that regulate genome accessibility by modulating histone acetylation status[37, 54]. Therefore, NAD levels are crucial for the function of Sir2. In our study, we detect a shortage of NAD in CTPS mutant cells, and not all NAD precursors can rescue non-quiescence states. We find that only methods that promote Sir2 functions, such as supplying nicotinic acid and overexpression of Sir2, can restore the cell fate of CTPS mutant to a quiescence state, indicating that Sir2 plays a key role in cytoophidium related quiescence state behind NAD levels.

As a downstream of Sir2, abnormally rDNA activation is observed in CTPS mutant, which is a determinant of cell fate [36, 55, 56]. Studies have shown that histone deacetylation[57] and rDNA condensation[58, 59] play a role in cell quiescence. Based on our results of low NAD levels and high Bna5 cytoophidium percentage in CTPS mutants, we believe that the Bna5-Glt1 cytoophidium complex restricts NAD supply and leads to Sir2 dysfunction, indicating that the cross-pathway regulatory mechanism of cytoophidium interactions ultimately leads to different cell fates. Therefore, we link cytoophidia with cell fate through Sir2 mediated approach. In short, our results indicate that metabolism can be regulated across pathways in a cytoophidium complex related way, thereby resonating with cell fates.

### Metabolic cell biology

Enzymes have multiple levels of regulations to control their activity, but there are few reports on enzyme regulation at the cellular biology level. In our previous studies, we have demonstrated that enzymes forming micron-level cytoophidia can regulate their own activity [14, 60–66]. In this study, we find a negative correlation between the formation of BNA pathway member Bna5 cytoophidium and NAD levels. Bna5 seems to be another example of how cytoophidium formation regulates its own enzymatic activity. In addition, we find that cytoophidium interactions can regulate metabolism that is not directly related to their own reactions.

CTPS cytoophidia and Glt1 cytoophidia can mediate NAD biosynthesis in opposite directions with Bna5 cytoophidium involved, which is a novel ectopic metabolic regulation system. Previous screening of cytoophidia in budding yeast showed that cytoophidium forming enzymes are more likely to be located at metabolic shunts[28]. Due to the communication role of enzymes at the shunt between pathways, their cytoophidia can serve as a physical means of multi pathway interactions. In this study, we demonstrate the inter pathway interactions between BNA, amino acid biosynthesis, and nucleotide biosynthesis through cytoophidium interactions. Overall, our research findings indicate the existence of metabolic cooperation at the cellular biology scale, bringing one of the first peaks in metabolic cell biology.

## MATERIALS AND METHODS

### Yeast strains and plasmids

All strains used in this study are derived from Saccharomyces cerevisiae S288c (BY4741) (**Supplementary Table S1**). All endogenous genomic tagging and plasmid transformation are done by TE/LiAc/PEG based transformation. Briefly, for endogenous genomic tagging, primers with 50-60bp homologous arms that anneal to the target genes are used to amplify DNA fragments with fluorescent protein (FP) tags. These DNA fragments will then be guided by the homologous arms to replace the stop codons or other specific regions of the target genes for fluorescent tagging. For the mutagenesis of genes, certain ORF will first be cloned into template FP plasmids. Then forward primers 50-100bp upstream the mutagenesis site will be used for amplifications and following transformations. The primers and plasmids used are listed in **Supplementary Tables S2 and S3**, respectively. The primers, template plasmids are all synthesized by Azenta Life Sciences, Inc (Suzhou, China).

### Yeast culturing, media and chemical treatments

All endogenous genomic tagged strains are cultured with liquid YPD (1% yeast extract, 2% peptone, 2% glucose) at 30℃, 250 rpm for certain days, 1 day for diauxic phase and 5 days for stationary phase. Nicotinic acid (NA), nicotinamide (NAM), cytosine are all dissolved in YPD at 16mM (3.2x), 25mM (5x) and 60mM(60x) respectively and diluted to 5mM, 5mM and 1mM as final concentration before culturing starts. The reagents and materials used are listed in **Supplementary Tables S4 and S5**, respectively.

### Confocal imaging and live cell imaging

For fixed samples, formalin are added to certain cultures to make a final 4% concentration and incubated at 30℃, 250rpm for 10 minutes. Centrifuged the samples at 12000rpm to discard supernatant. Washed with 1x PBS once and then resuspend in 20uL 1x PBS for imaging. Samples are mixed with calcofluor white (0.0075% w/v) and low-melting point agarose (LMA, 1.2%) at a ratio of 2.5:1:1 and imaged by ZEISS LSM 980 AiryScan2 with a 63x oil objective.

Short-term live cell imaging of starvation induction: Cells are first incubated 6-8 hours in 2mL YPD from OD600=0.05 to 0.8 them centrifuged at 4000g for 1 minutes. Then the pellet is washed by 500μL 5-day YPD prepared previously (filtered by a 0.22μm filter, Millipore) and resuspend in 20uL 5-day YPD. Drip 5uL of the culture onto a glass bottom dish (MatTek) and 80uL pre-melted 1.5% LMA in 5-day YPD.

Stir to mix and solidify the LMA mix in 30℃ for around 5min. Add 1mL 5-day YPD in the dish finally. Transfer the dish to ZEISS LSM 980 AiryScan2 confocal microscope with a 63x oil objective at 30℃. Images are taken every 5 minutes for 90 minutes in total.

Long-term live cell imaging in YPD: Cells are first incubated 6-8 hours in 2mL YPD from OD600=0.05 to 0.8 them centrifuged at 4000g for 1 minutes. Concentrate the cells to 20uL. Use 1mL supernatant to dissolve 15mg LMA in 60℃. Drip 5uL of the cell onto a glass bottom dish mentioned above. Mix the cells with 80uL pre-melted 1.5% LMA in YPD just separated and solidify the LMA mix in 30℃ for around 5min. Add the remaining supernatant in the dish and seal with parafilm finally. Transfer the dish to ZEISS Cell Observer SD spinning disk confocal microscope with a 63x oil objective at 30℃. Images are taken every 15 minutes for 3 days in total.

### Fractionation of quiescent cells and non-quiescent cells

Percoll gradient (Percoll:1.5M NaCl = 8:1 in volume) is prepared by centrifuging at 19300g in a Beckman super centrifuging tube by Optima XPN-100 centrifuge with a SW41 rotor. 500uL 5-day live cells or fixed cells are harvested, washed by 1x PBS and concentrated to 200uL in 1xPBS. Lay the 200uL cells on the top of the gradient, seal with parafilm. Centrifuge the cells at 400g for 1hour in Eppendorf 5810R centrifugate with swinging buckets. Separate the cells with pipette and wash with 5mL 1x PBS twice. For live cell experiments, e.g. transcriptome, fractionated cells will be resuspended in 5-day YPD for 2 hour, 30℃, 250 rpm.

### Chronological life span assay

Stains are first diluted to OD600= 0.05 and cultured in YPD for 3 days. At the third day, dilute the culture 1000 times and spread 10uL of diluted culture onto YPD agar (2%) plates. Repeat this dilution every other day and count the colonies formed on the last plate. Take the colony number of the third day as the 100% survival rate and divide the colony numbers by which of the third day. Plot the curve of chronological life span with Prism GraphPad 10 (v10.3.2).

### Spot assay

Stains are first diluted to OD600= 0.05 and cultured in YPD for 1 day or 5days. Adjust OD600 to 0.1 by water, dilute it 10 times for 3 times and get the dilutions of OD600= 0.1, 0.01, 0.001, 0.0001 respectively. Drip 2.5 μL of each dilution onto YPD agar plates and grow for 2 days. Images are taken by a Tanon-1000 CCD camera.

### ROS detection assay

1uL of 40mM of DHE (Dihydroethidium, SparkJade) is added to 1mL of culture at certain stages (5 days or 9 days) in YC medium. Incubate at 30℃ for 1 hour, 250 rpm. Then wash with YC medium separated from certain stage cultures once. Resuspend in 20μL YC medium from separated from certain stage cultures. Mix 2.5 μL of cells with 1μL of 1.2% LMA on the slide for further confocal imaging by ZEISS LSM 980 AiryScan2. An mCherry channel is used for DHE excitation and emission.

### Replicative age analysis

Cells are fixed and collected as described in *Confocal imaging and live cell imaging.* Mix 2.5μL of cells with 1μL of 1.2% LMA and 1μL of calcofluor white (0.0075% w/v) for imaging. Cells with more than 2 scars are recorded. Percentage s of cells with more than 2 scars are calculated by dividing the total number of cells checked.

### Transcriptomic profiling

3 OD of CTPS mutant, UGPA control, CTPS OE, upper CTPS mutant and lower CTPS mutant cells are collect at 8 hours or 5 days of culturing in YPD or YPDK (with 0.2% G418, for CTPS OE and UGPA group). Cells are transferred into 500μL TriZol reagent (Transgene) with ZrO beads and lysed in a homogenizer shaking at 75Hz for 1 minute for 6 cycles with 1.5 minute break on ice between cycles. The cell lysates are sent to the Next Gen Sequencing Department of Azenta Life Sciences, Inc (Suzhou, China) for mRNA library constructing, sequencing and data analysis. An Illumina HiSeq is used for sequencing according to manufacturer’s instructions. Differential expression analysis used the DESeq2 Bioconductor package, a model based on the negative binomial distribution. the estimates of dispersion and logarithmic fold changes incorporate data-driven prior distributions, P-value of genes were set <=0.05 to detect differential expressed ones.

Differentially expressed genes of CTPS mutant and CTPS OE are combined for discovering the intersection genes. These genes both differentially expressed in two strains are used for GO and KEGG pathway analysis by using YeastEnricher[67].

### Metabolomic profiling

3 OD of CTPS mutant and UGPA control cells are collected at 5 days cultured in YPD. Cells are directly frozen in liquid nitrogen and sent to Obio Tech, Inc for high-throughput targeted metabolomics, using a H650 kit (Applied Protein Technology Co., Ltd.). For the liquid chromatography-mass spectrometry (LC-MS) analysis, samples were dissolved in 100 μL of a 1:1 (v/v) acetonitrile/water solvent and centrifuged at 14,000 g for 15 minutes at 4 °C. The resulting supernatant was then injected for analysis. This was conducted using a 1290 Infinity LC (Agilent Technologies) coupled with a QTRAP MS 6500+ (AB Sciex) at Obio Tech, Inc. The analytes were separated using HILIC (Waters UPLC BEH Amide column, 2.1 mm × 100 mm, 1.7 μm) and C18 columns (Waters UPLC BEH C18, 2.1 × 100 mm, 1.7 μm). Quality control (QC) samples were included in the sample queue to assess the stability and repeatability of the system.

Data are analysed by MultiQuant and Analyst. Principal component analysis (PCA) is used for data clustering. One-way ANOVA is performed for comparisons.

### Data analysis of confocal images

Statistics of confocal images are done all manually with more than 300 cells per biology replicant. Data analyses and plotting are done by GraphPad Prism v10.2.3. Two-way ANOVA is used for significance analysis. Colocalization profiling is presented by ZEISS Zen lite 3.2.

## ACKNOWLEDGMENTS

We thank all the members of Ji-Long Liu Lab for supporting and instructing. We thank the Molecular Imaging Core Facility and Molecular and Cell Biology Core Facility at the School of Life Science and Technology, ShanghaiTech University for providing technical support. This work was supported by the grants from the Ministry of Science and Technology of China (No. 2021YFA0804700), National Natural Science Foundation of China (Grant Nos. 32370744 and 32350710195), and UK Medical Research Council (Grant Nos. MC_UU_12021/3 and MC_U137788471) for grants to J. L. L.

---

**Supplementary Table S1.**
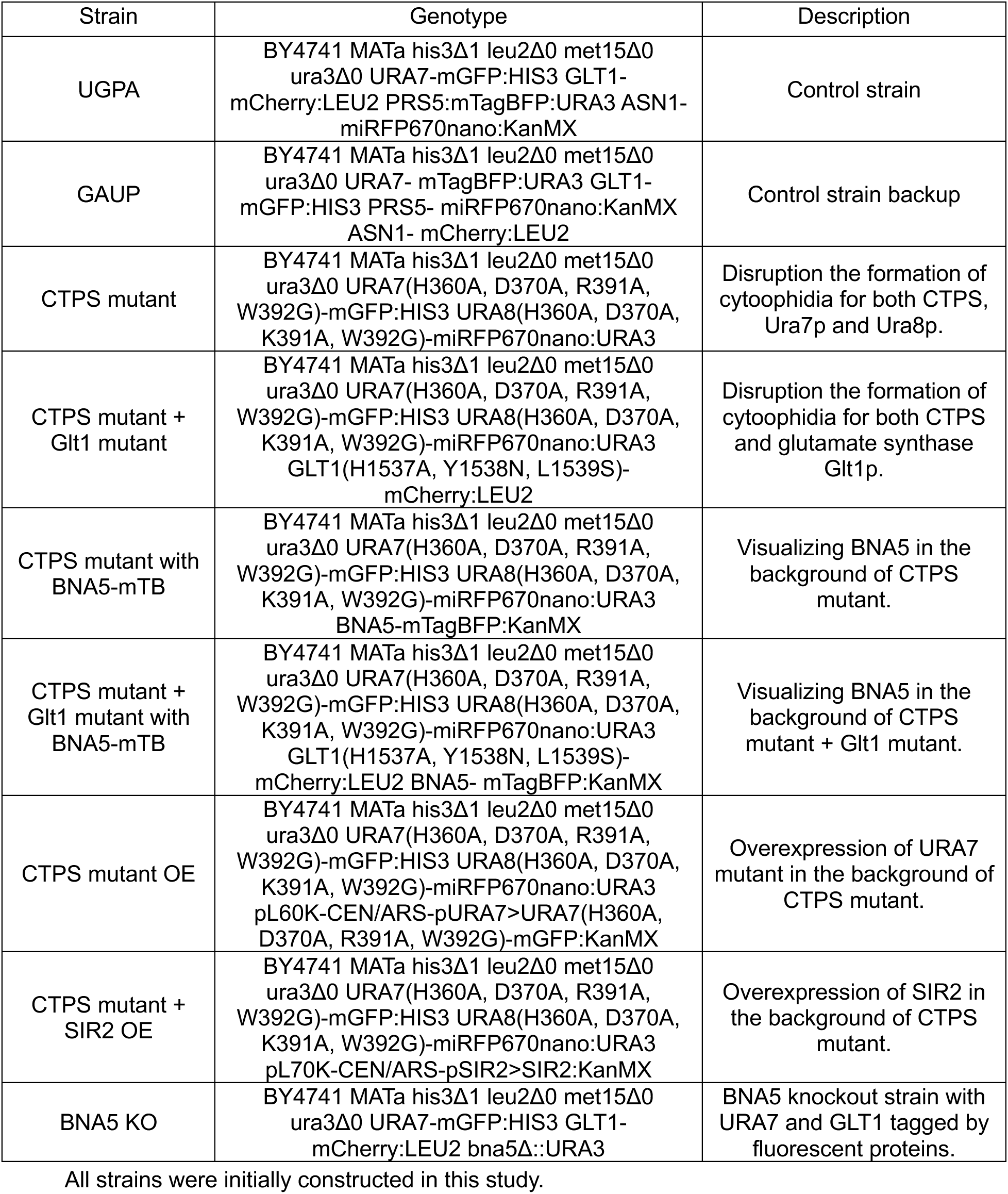
The strains used in this study.

**Supplementary Table S2.**
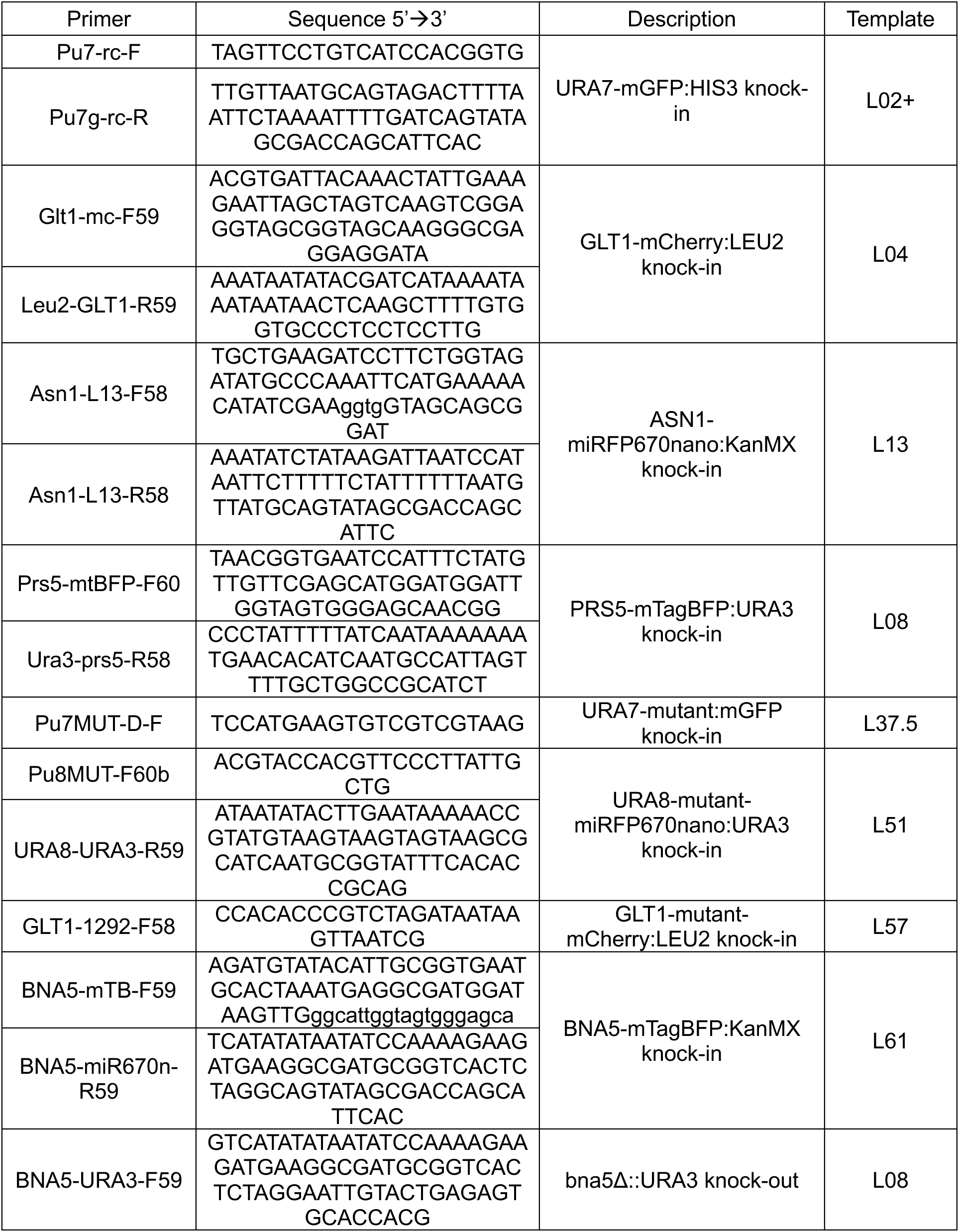
The primers used in this study.

**Supplementary Table S3.** The plasmids used in this study.

| Plasmids | Genotype | Description |
| --- | --- | --- |
| L02+ | pL02-URA7-mGFP(S65T, A206K)-ADH1ter-pTEF>HIS3-TEFter-AmpR | Used for mGFP tagging in genome. |
| L04 | pL04-CEN/ARS-mCherry-ADH1ter<br>pLEU2>LEU2-AmpR | Used for mCherry tagging in genome. |
| L08 | pL08-CEN/ARS-mTagBFP-ADH1ter<br>pURA3>URA3-AmpR | Used for mTagBFP tagging in genome. Used for gene replacement by URA3 as well. |
| L13 | pL13-CEN/ARS-pGAP>LifeAct-miRFP670nano-ADH1ter-KanMX-AmpR | Used for miRFP670nano tagging in genome. |
| L37.5 | pL37.5-URA7(H360A, D370A, R391A, W392G)-mGFP-ADH1ter-pHIS3>HIS3-AmpR | Used for URA7 endogenous gene mutation. |
| L51 | pL51-URA8(H360A, D370A, k391A, W392G)-miRFP670nano-ADH1ter-pURA3>URA3-AmpR | Used for URA8 endogenous gene mutation. |
| L57 | pL57-CEN/ARS-GLT1(H1537A, Y1538N, L1539S)-mCherry:LEU2-AmpR | Used for Glt1 endogenous gene mutation. |
| L60K | pL60K-CEN/ARS-pURA7>URA7(H360A, D370A, R391A, W392G)-mGFP:KanMX-AmpR | Used for URA7 mutant overexpression |
| L61 | pL61-CEN/ARS-mTagBFP-ADH1ter<br>KanMX-AmpR | Used for mTagBFP tagging in genome, but with a KanMX selection |
| L70K | pL70K-CEN/ARS-pSIR2>SIR2:KanMX-AmpR | Used for SIR2 overexpression |

All plasmids were initially synthesized in this study.

**Supplementary Table S4.** Reagents.

| Reagents | Brand | Cat Number | Description |
| --- | --- | --- | --- |
| Phanta Max Super-Fidelity DNA Polymerase | Vazyme | P505 | PCR |
| 2X MultiF Seamless Assembly Mix | ABclonal Technology | RK21020 | Gibson assembly |
| Lithium acetate dihydrate | Macklin | L812345 | Yeast transformation |
| PEG4000 | Meilunbio | MB2587 | Yeast transformation |
| TE (Tris/EDTA buffer) | Beyotime Biotechnology | ST725 | Yeast transformation |
| ssDNA | Sigma-Aldrich | D9156 | Yeast transformation |
| DpnI | ABclonal Technology | RK21109 | Template digestion |
| PCR cleanup kit | UElandy | UE-PCR-250 | PCR cleanup |
| Nicotinic acid | Accela ChemBio | SY011111 | Chemical addition |
| Nicotinamide | Accela ChemBio | SY024804 | Chemical addition |
| Cytosine | Accela ChemBio | SY001643 | Chemical addition |
| Yeast Extract | Oxoid | LP0021B | Yeast culture |
| Peptone | Sigma-Aldrich | 82962 | Yeast culture |
| Glucose | Solarbio | G8150 | Yeast culture |
| YNB w/o AA or (NH<sub>4</sub>)<sub>2</sub>SO<sub>4</sub> | BBI | A600505 | Yeast culture |
| Succinic acid | Collins | P2819863 | Yeast culture |
| Ammonium sulfate | Sinopharm Chemical Reagent | 10002918 | Yeast culture |
| Tyrosine | Sigma-Aldrich | T8566 | Yeast culture |
| Methionine | Sigma-Aldrich | M5308 | Yeast culture |
| Arginine | Sigma-Aldrich | A5006 | Yeast culture |
| Alanine | Sigma-Aldrich | A7469 | Yeast culture |
| Serine | Sigma-Aldrich | S4311 | Yeast culture |
| Valine | Sigma-Aldrich | V0513 | Yeast culture |
| Threonine | Sigma-Aldrich | T8841 | Yeast culture |
| Isoleucine | Sigma-Aldrich | I7403 | Yeast culture |
| Glycine | Sigma-Aldrich | G8790 | Yeast culture |
| Phenylalanine | Sigma-Aldrich | P5482 | Yeast culture |
| Cysteine | Sigma-Aldrich | C7352 | Yeast culture |
| Aspartic acid | Sigma-Aldrich | 17219 | Yeast culture |
| Glutamic acid | Sigma-Aldrich | G1626 | Yeast culture |
| Proline | Sigma-Aldrich | P5609 | Yeast culture |
| Tryptophane | Sigma-Aldrich | T8941 | Yeast culture |
| Leucine | Sigma-Aldrich | L8912 | Yeast culture |
| Histidine | Sigma-Aldrich | H5659 | Yeast culture |
| Uracil | Sigma-Aldrich | U1128 | Yeast culture |
| Lysine | Sigma-Aldrich | L8662 | Yeast culture |
| G418 | GPC | AK108 | Yeast culture |
| Sodium hydroxide | Sinopharm Chemical Reagent | 10019718 | Yeast culture |
| Agar | Abcone | A42307 | Yeast culture |
| Fluorescent Brightener 28<br>disodium salt solution<br>(CFW) | Sigma-Aldrich | 910090 | Imaging |
| Low gelling point agarose | Sigma-Aldrich | A4018 | Imaging |
| Formalin | Sigma-Aldrich | F8775 | Imaging |
| Dihydroethidium | SparkJade | SJ-MD0019 | Imaging |
| Percoll | Yeasen | 40501ES60 | Cell separation |
| Sodium chloride | Sinopharm Chemical Reagent | 10019318 | Cell separation |
| 20x PBS buffer | Sangon Biotech | B548117 | Buffer |
| TransZol Up | Transgen | ET111-01 | RNA extraction |

**Supplementary Table S5.** Materials.

| Materials | Brand | Cat Number | Description |
| --- | --- | --- | --- |
| 9cm petri dish | PULLEN | PX02003 | Yeast culture |
| 35mm Glass Bottom Dish | MatTek | P35G-1.5-10-C | Imaging |
| Cover slip | Brand | 470050 | Imaging |
| Adhesion Slide | Titan | SWBP-Z0003 | Imaging |
| Glass beads | Sigma-Aldrich | G8772 | RNA extraction |
| Syringe Filter, 0.22 µm | Millipore | SLGPR33RB | Sterilization |
| 13.2mL Open-Top Ultra-Clear Tube | Beckman Coulter | 344059 | Ultracentrifugation |

**Figure S1.**
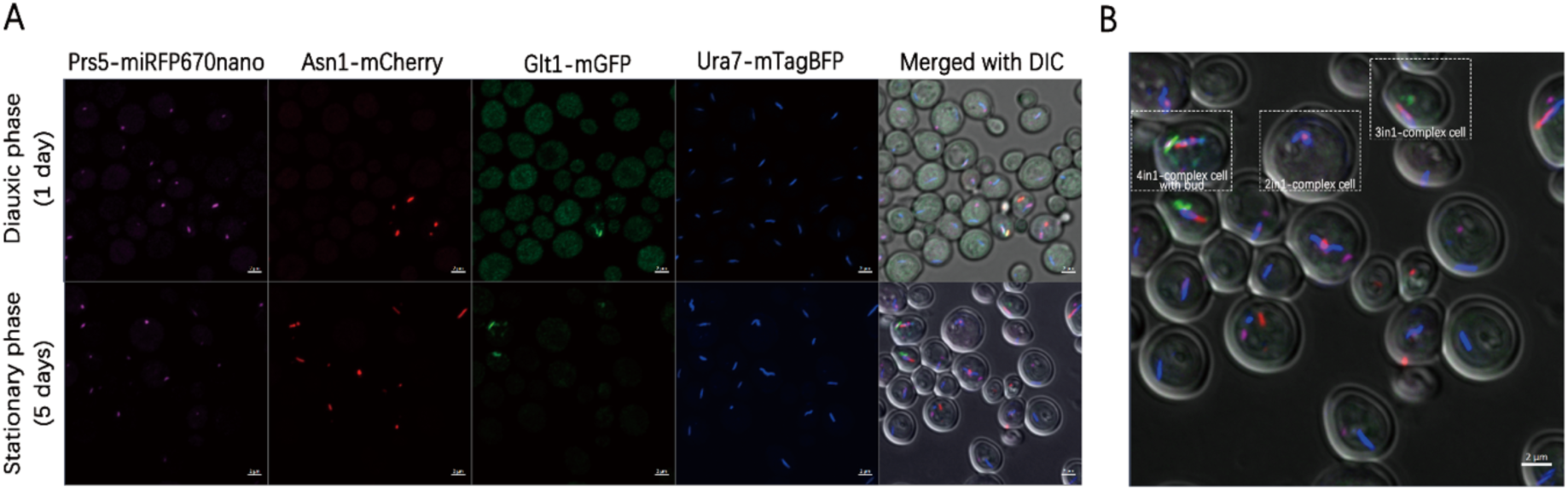
The formation of cytoophidium complex is independent of fluorescent protein tags. **A.** The formation of cytoophidia and complexes during different growth stages. Purple represents Prs5-miRFP670nano, red represents Asn1-mCherry, green represents Glt1-mGFP, blue represents Ura7-mTagBFP. Scale bar = 2 μm. **B.** Zoom-in of stationary phase cells in A, and cells with different complexes are labelled by squares. Scale bar = 2 μm.

**Figure S2.**
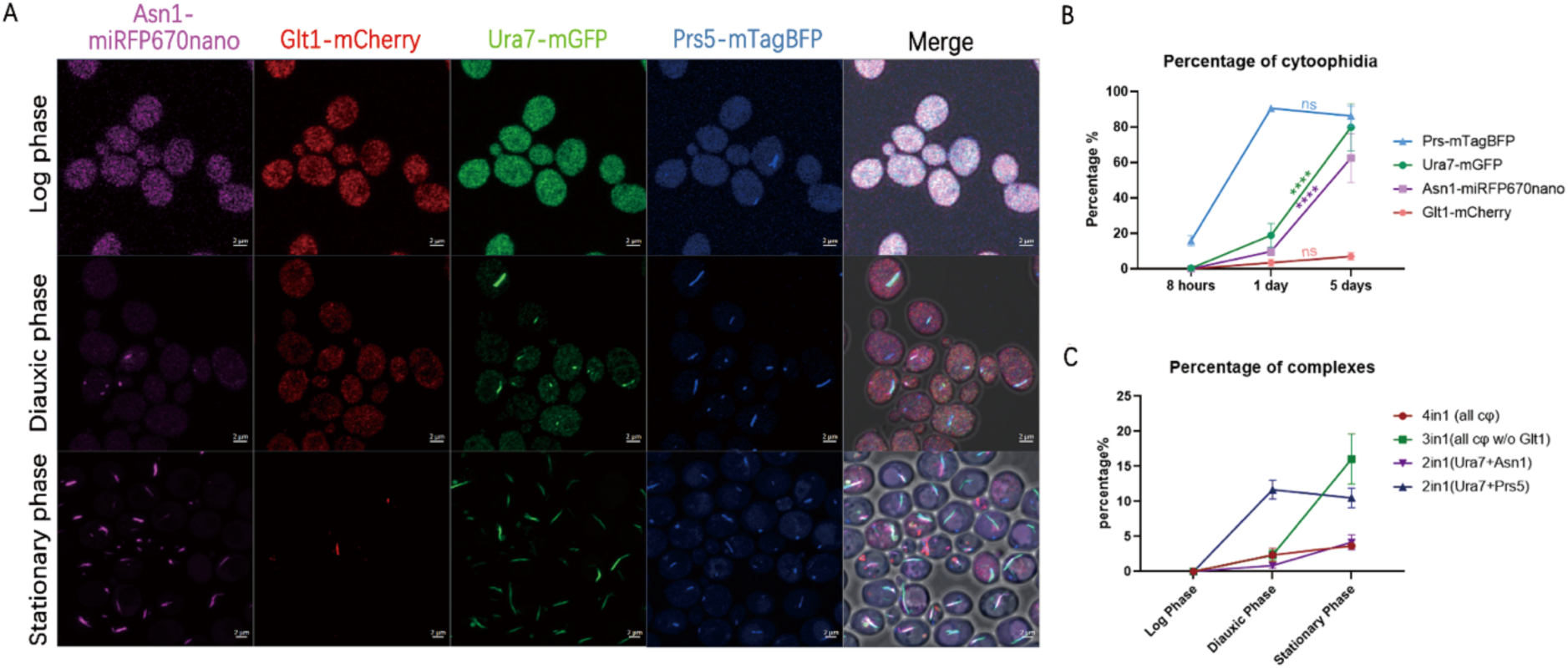
Cytoophidium abundance and complex abundance during cultivation. **A.** The formation of cytoophidia and complexes during different growth stages. Purple represents Asn1-miRFP670nano, red represents Glt1-mCherry, green represents Ura7-mGFP, blue represents Prs5-mTagBFP. Scale bar = 2 μm. **B.** Quantification of the abundance of different cytoophidia during different growth stages. **C.** Quantification of the abundance of cytoophidium complexes with different compositions.

**Figure S3.**
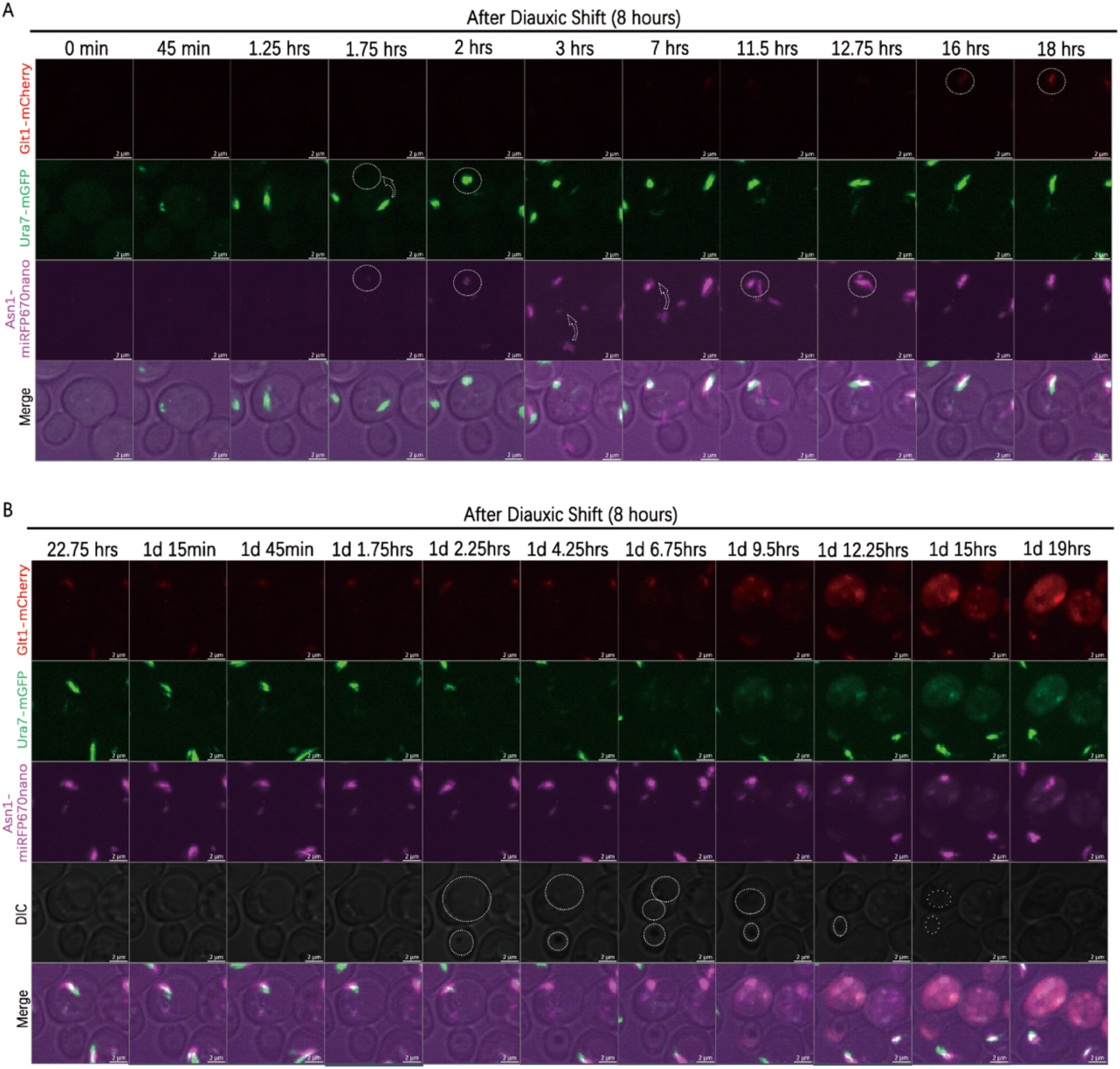
Live imaging of Ura7-Glt1 cytoophidium interactions. **A.** The sequence of cytoophidium formation during normal culturing includes Ura7, Asn1, and Glt1 cytoophidium. Circles indicate areas of cytoophidium occurrence. Arrows indicate the movement of cytoophidia. **B.** Sequence of cytoophidium remodeling and cell shrinkage. Circles indicate the size of vacuoles in both mother and daughter cells. Scale bar = 2 μm.

**Figure S4.**
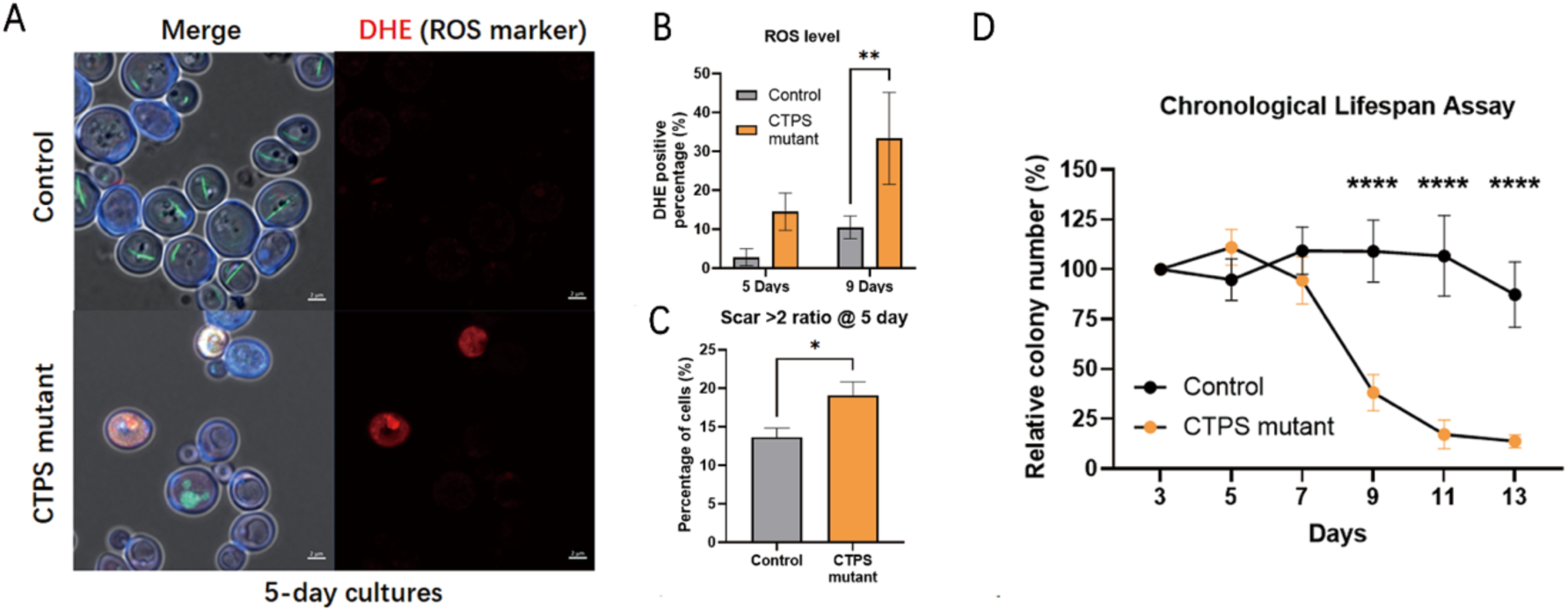
The non-quiescent phenotype of CTPS mutant. **A.** Confocal images of ROS marker DHE (red) in Control strain and CTPS mutant at stationary phase (5 days). Ura7-mGFP (green), CFW (cell wall, blue), DIC (white). **B.** ROS level plot of strains in A at stationary phase (5 days and 9 days). **C.** The percentage of cells with 2 or more scars in the control strain and CTPS mutant at stationary phase (5 days). D. The chronological survival rate of the control strain and CTPS mutant from 3 days to 13 days. * means p-value < 0.05; **, p-value < 0.01; *** p-value < 0.001; ****, p-value < 0.0001; and ns or not labelled mean no significance in the statistical chart.

**Figure S5.**
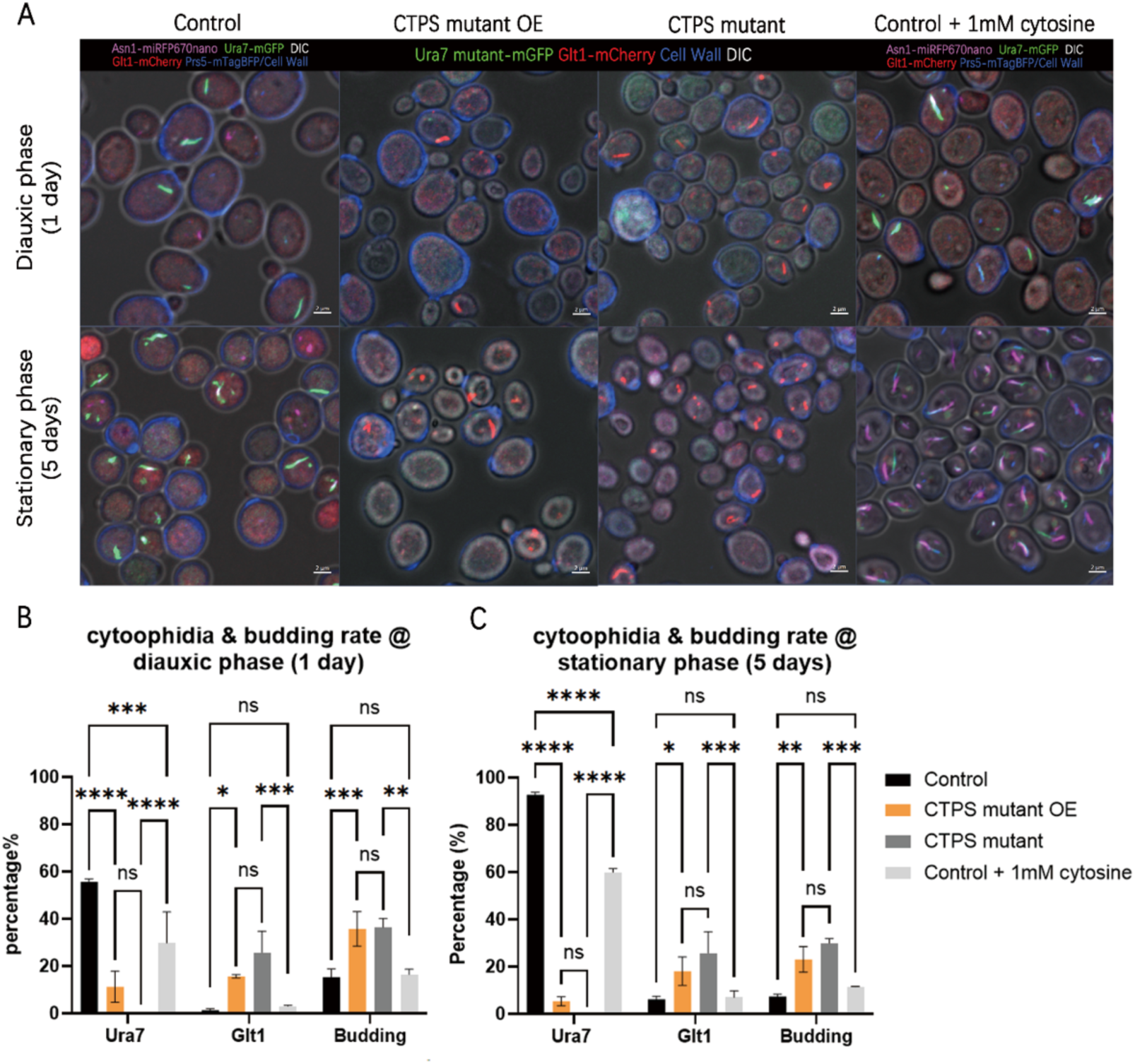
The non-quiescent phenotype is independent of the expression level or enzymatic activity of CTPS. **A.** Confocal images of control strain (UGPA), CTPS mutant (Ura7^mutant^-mGFP, Ura8^mutant^-miRFP670nano, Glt1-mCherry), CTPS mutant OE (a URA7^mutant^-mGFP overexpression plasmid within CTPS mutant strain) and Control treated with 1 mM cytosine at diauxic phase (1 day) and stationary phase (5 days). For the control strain with complex, Ura7 signal (green), Glt1 signal (red), Prs5-mTagBFP /CFW (blue), Asn1-miRFP670nano and DIC (white) are shown. For the mutants, only Ura7 signal (green), Glt1 signal (red), CFW (cell wall, blue) and DIC (white) are shown. Scale bar, 2μm. **B,C.** Plots showing the percentages of Ura7 cytoophidium, Glt1 cytoophidium and budding rate in strains mentioned in A at diauxic phase (1 day, B) and stationary phase (5 days, C). * means p-value < 0.05; **, p-value < 0.01; ***, p-value < 0.001; ****, p-value < 0.0001; and ns or not labelled mean no significance in the statistical chart.

**Figure S6.**
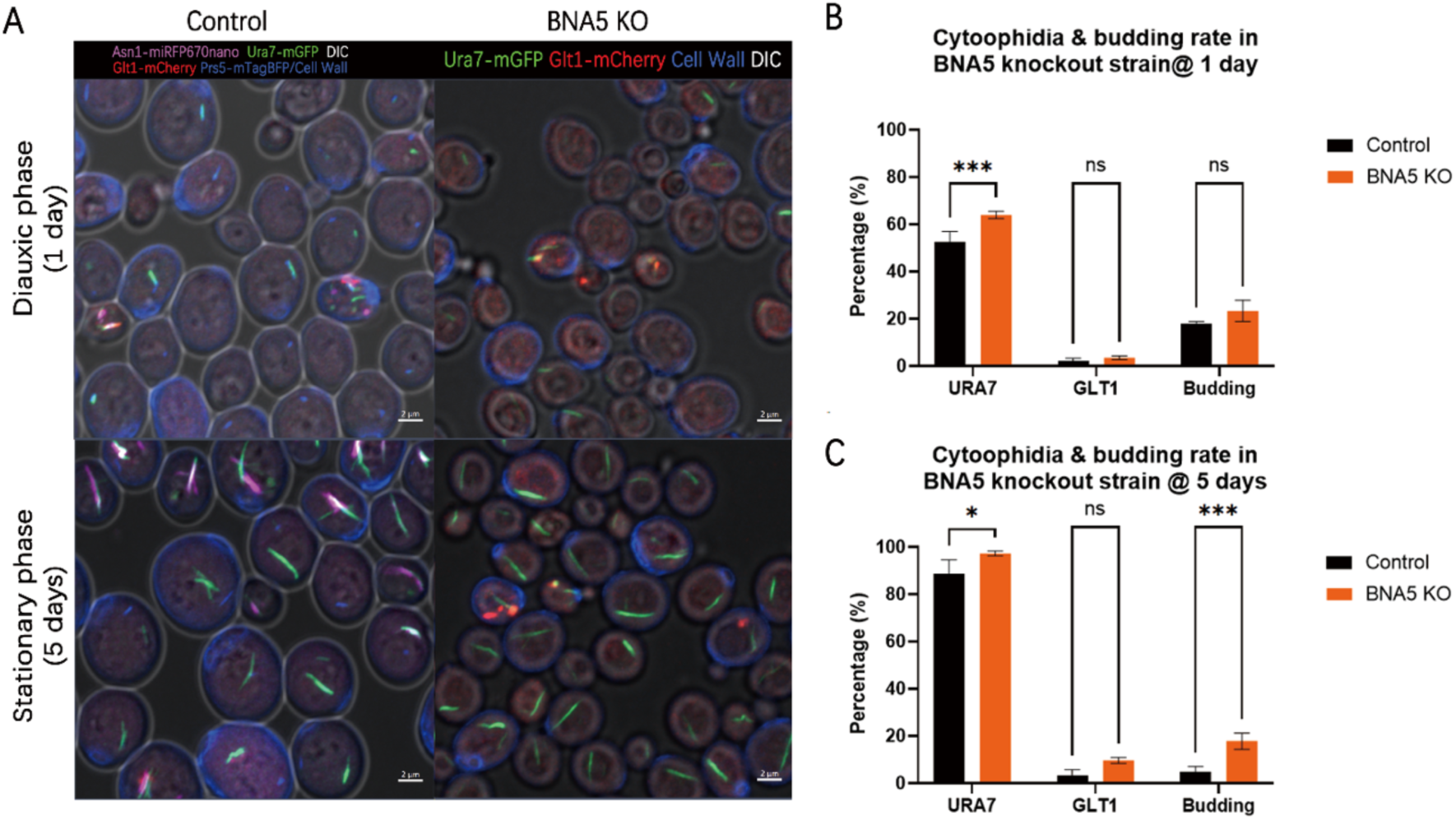
Knockout of BNA5 can lead to a higher budding rate at stationary phase. **A.** Confocal images of control strain (UGPA) and BNA5 KO (with URA7-mGFP & Glt1-mCherry) at diauxic phase (1 day) and stationary phase (5 days). **B,C.** Quantification of URA7, GLT1 cytoophidia percentages among cells and the budding rate of the strains mentioned in A at 1 day (B) and 5 days (C). Scale bar = 2 μm. * means p-value < 0.05; **, p-value < 0.01; ***, p-value < 0.001; ****, p-value < 0.0001; and ns or not labelled mean no significance in the statistical chart.

## Notes

### Competing Interest Statement

The authors have declared no competing interest.

